# Epileptic phenotypes in *slc13a5* loss-of-function zebrafish are rescued by blocking NMDA receptor signaling

**DOI:** 10.1101/2024.01.15.575806

**Authors:** Deepika Dogra, Van Anh Phan, Cezar Gavrilovici, Nadia DiMarzo, Kingsley Ibhazehiebo, Deborah M. Kurrasch

**Author notes:** **Correspondence** Deborah M. Kurrasch, University of Calgary, 3330 Hospital Drive NW, HSC 2215, Calgary, Alberta T2N 4N1.

## Abstract

*SLC13A5* encodes a citrate transporter highly expressed in the brain important for regulating intra- and extracellular citrate levels. Mutations in this gene cause a rare infantile epilepsy characterized by lifelong seizures, developmental delays, behavioral deficits, poor motor progression, and language impairments. SLC13A5 individuals respond poorly to treatment options; yet drug discovery programs are limited due to a paucity of animal models that phenocopy human symptoms. Here, we used CRISPR/Cas9 to create loss-of-function mutations in *slc13a5a* and *slc13a5b*, the zebrafish paralogs to human *SLC13A5*. *slc13a5* mutant larvae showed cognitive dysfunction and sleep disturbances, consistent with SLC13A5 individuals. These mutants also exhibited fewer neurons and a concomitant increase in apoptosis across the optic tectum, a region important for sensory processing. *slc13a5* mutants displayed hallmark features of epilepsy, including an imbalance in glutamatergic and GABAergic excitatory-inhibitory gene expression, disrupted neurometabolism, and neuronal hyperexcitation as measured *in vivo* by extracellular field recordings and live calcium imaging. Mechanistically, we tested the involvement of NMDA signaling in *slc13a5* mutant epilepsy-like phenotypes. Slc13a5 protein co-localizes with excitatory NMDA receptors in wild-type zebrafish and blocking NMDA receptors in *slc13a5* mutant larvae rescued bioenergetics, hyperexcitable calcium events, and behavioral defects. These data provide empirical evidence in support of the hypothesis that excess extracellular citrate over-chelates the ions needed to regulate NMDA receptor function, leading to sustained channel opening and an exaggerated excitatory response that manifests as seizures. These data show the utility of *slc13a5* mutant zebrafish for studying SLC13A5 epilepsy and open new avenues for drug discovery.

## Introduction

SLC13A5 is a sodium-coupled citrate transporter (NaCT), encoded by the *SLC13A5* gene in mammals and expressed in the plasma membrane of liver, testis, bone and neural cells (1–4). In the brain, the citrate transporter is expressed predominantly in neurons where it mediates the uptake of circulating and astrocyte-released citrate. Citrate plays a crucial role in lipid and cholesterol biosynthesis, as well as in the regulation of energy expenditure (5). Mutations in *SLC13A5* cause developmental epileptic encephalopathy 25 (DEE25) characterized by prolonged and frequent seizures as early as the first day of life (6–10). Other associated comorbidities include developmental delays, teeth hypoplasia, cognitive impairments, sleeping difficulties and slow motor progress (11–13). Its *Drosophila* homolog, *Indy,* initially was associated with longevity (14) but upon re-examination was found to exhibit a loss of glutamatergic neurons and an overlooked “bang-induced” seizure-like phenotype (15). *Slc13a5* knockout mice are small compared to wild-type controls and display metabolic phenotypes such as protection against obesity and insulin resistance (16). Although originally thought to lack any epilepsy-related phenotypes, more recently *Slc13a5-* null animals were found to display neural hyperexcitability but no obvious behavioral or histological abnormalities (17), questioning the utility of mice in modeling the human SLC13A5 syndrome given the strong seizures experienced. Recently, Dirckx et al. reported an osteoblast metabolic pathway dysregulated in *Slc13a5^−/−^*mice, revealing insights into the teeth and bone deformations observed in humans (18).

Individuals suffering from SLC13A5 epilepsy harbor either homozygous or compound heterozygous *SLC13A5* pathogenic variants. Anti-seizure medications such as stiripentol or a combination of acetazolamide and valproic acid reduce seizure frequency in some SLC13A5 individuals; however most children are refractory and complete seizure freedom is never attained even in those that do respond to anti-seizure medications (13). *In vitro* functional studies on several *SLC13A5* mutations have revealed disrupted transporter activity pointing towards a loss of function (6, 19), but the molecular mechanisms contributing to seizures are still poorly understood.

Zebrafish are a powerful system for studying epilepsy with several genetic and pharmacologically induced models recapitulating brain epileptiform activity (20–23). Moreover, zebrafish larvae are a useful vertebrate model for understanding the etiology of neurological disorders given the range of behavioral assessments, genetic manipulations, and high-resolution measures of brain activity. To explore epilepsy-related phenotypes due to the loss of *SLC13A5* and study the molecular mechanisms behind this disorder, we generated the genetic mutants of both zebrafish *SLC13A5* paralogs, *slc13a5a* (referred to as *5a*) and *slc13a5b* (referred to as *5b*). Here, we characterized the phenotypes of all three mutants: *5a^−/−^* singly, *5b^−/−^*singly, and *5a^−/−^;5b^−/−^* double mutants. Additionally, we studied the role of NMDA receptor signaling in *SLC13A5* epileptic phenotypes. Of note: to simplify the language throughout this text, hereafter we use “*slc13a5^−/−^*” or “*slc13a5* mutants” to refer to any one of these three mutant strains. The exact mutant analyzed is defined in the figures.

## Results

### *slc13a5* is expressed in the brain and *slc13a5*^−/−^ larvae display a smaller midbrain and increased mortality

Due to an ancestral gene duplication event, zebrafish possess two paralogs of *SLC13A5*: *5a* and *5b*. We performed in situ hybridization for *5a* and *5b* expression and found that both paralogs were highly expressed in the zebrafish brain. Widespread expression was observed across the central nervous system during the early developmental stages of 25 hours post fertilization (hpf) and 3 days post fertilization (dpf); however, this expression became primarily restricted to the midbrain (MB) by 5 dpf (Fig. 1A-B”, Fig. S1A, B). To examine the neural cell types expressing Slc13a5, we performed co-immunostaining for Slc13a5 and either HuC/D (neuron marker) or Gfap (astrocyte marker) and observed that Slc13a5 is expressed both in the neurons and astrocytes in the MB region of 3 dpf zebrafish larvae (Fig. 1C and D).

**Fig. 1.**
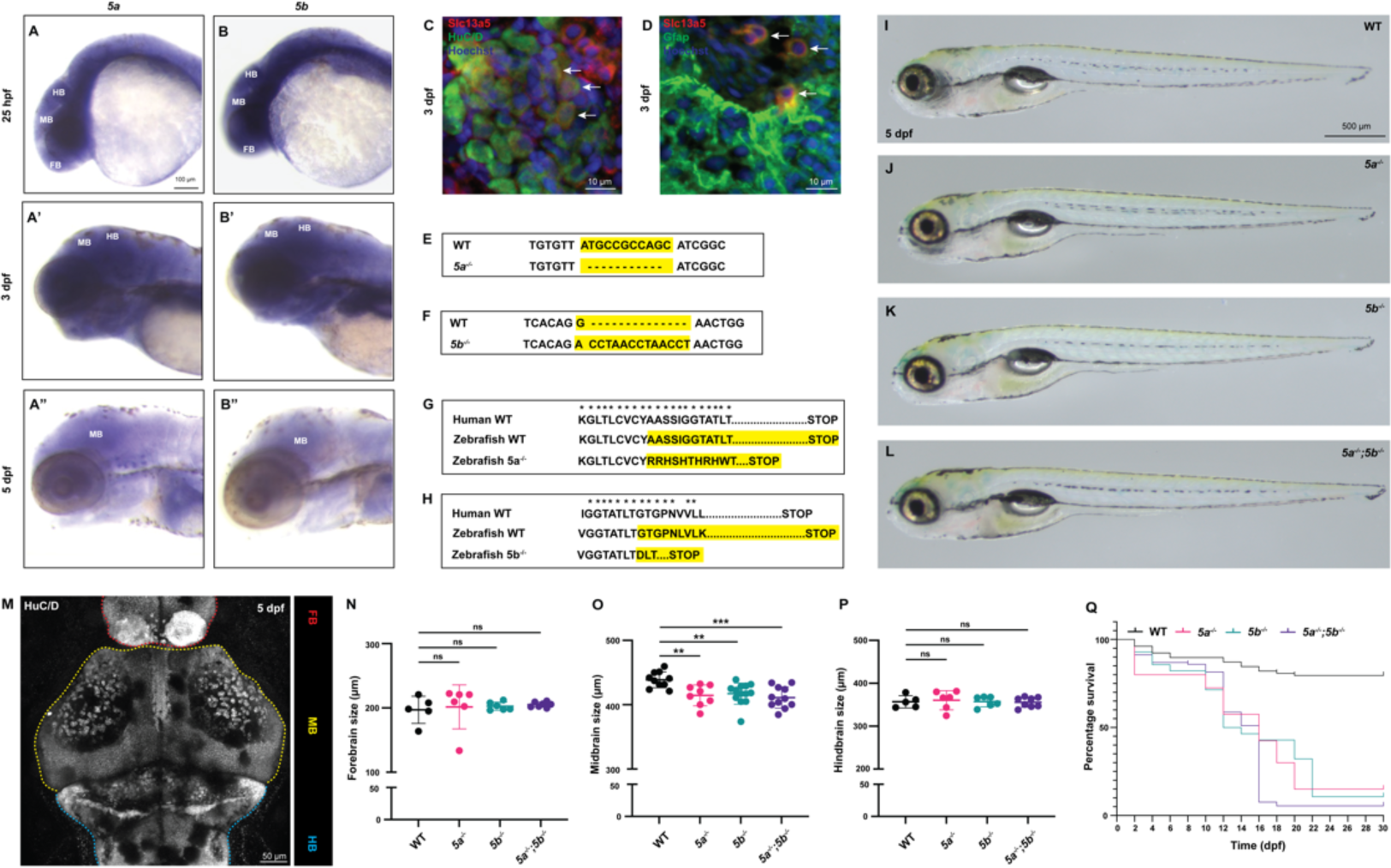
Expression analysis of *slc13a5* zebrafish paralogs (*5a* and *5b*) and generation of zebrafish *slc13a5* mutants using CRISPR/Cas9 technique. (A-B”) In situ hybridization for *5a* and *5b* expression at 25 hpf, 3 dpf and 5 dpf. *5a* and *5b* are expressed in all the brain regions during early development and their expression gets restricted to MB at later stages. (C) Section of 3 dpf WTs; α-Slc13a5, α-HuC/D, Hoechst. White arrows point to Slc13a5-expressing neurons (Slc13a5^+^/HuC/D^+^) in MB. (D) Section of 3 dpf WTs; α-Slc13a5, α-Gfap, Hoechst. White arrows point to Slc13a5-expressing astrocytes (Slc13a5^+^/Gfap^+^). (E) Nucleotide sequences of *5a* in zebrafish WT and *5a^−/−^*. Yellow highlighted regions indicate 11-nucleotide deletion due to mutation in *5a*. (F) Nucleotide sequences of *5b* in zebrafish WT and *5b^−/−^*. Yellow highlighted regions indicate one-nucleotide substitution followed by 13-nucleotide insertion due to mutation in *5b*. (G) Amino acid sequence of 5a in human orthologue, zebrafish WT and *5a^−/−^*. Asterisks indicate conserved amino acids between human and zebrafish. Yellow highlighted regions indicate affected amino acids and premature stop due to mutation in *5a*. (H) Amino acid sequence of 5b in human orthologue, zebrafish WT and *5b^−/−^*. Asterisks indicate conserved amino acids between human and zebrafish. Yellow highlighted regions indicate affected amino acids and premature stop due to mutation in *5b*. (I-L) WT and *slc13a5* mutant larvae at 5 dpf. No morphological differences were observed. WT, n=5; *5a^−/−^,* n=5; *5b^−/−^*, n=5; *5a^−/−^;5b^−/−^*, n=5. (M-P) Quantification of FB, MB and HB size at 5 dpf. MB size is significantly reduced in *slc13a5* mutants; however, FB and HB size remained unchanged. WT, n=5; *5a^−/−^,* n=6; *5b^−/−^*, n=6; *5a^−/−^;5b^−/−^*, n=7 for FB size measurement. WT, n=10; *5a^−/−^,* n=8; *5b^−/−^*, n=13; *5a^−/−^;5b^−/−^*, n=11 for MB size measurement. WT, n=5; *5a^−/−^,* n=6; *5b^−/−^*, n=6; *5a^−/−^;5b^−/−^*, n=8 for HB size measurement. (Q) Survival curve of WTs and *slc13a5* mutants. Very few *slc13a5* mutants survive until 30 dpf. WT, n=78; *5a^−/−^,* n=40; *5b^−/−^*, n=28; *5a^−/−^;5b^−/−^*, n=92. Data are Mean ± S.D., ns: no significant changes observed, **P ≤ 0.01, ***P ≤ 0.001-Unpaired t-test. hpf, hours post fertilization; dpf, days post fertilization; WT, wild-type siblings; *5a*, *slc13a5a*; *5b*, *slc13a5b*; FB, forebrain; MB, midbrain; HB, hindbrain.

Next, we injected CRISPRs targeting the fifth exons of *5a* and *5b* genes to generate mutant alleles, *5a^−/−^* and *5b^−/−^*. *5a^−/−^* allele mutations comprised an 11-nucleotide deletion and *5b^−/−^* allele mutations comprised one-nucleotide substitution followed by 13-nucleatide insertion, both predicted to encode truncated proteins due to a premature stop (Fig. 1E-H). These mutations were targeted in the sodium-binding site of NaCT, conserved among humans and zebrafish (Fig. 1G and H). Notably, several variants of SLC13A5 epilepsy have mutations in the sodium-binding site of NaCT, suggesting that sodium binding deficiency causes impaired citrate transport (7). To assess gene expression levels, we used quantitative PCR (qPCR) at 5 dpf and showed that *5a* transcript levels were significantly reduced in *5a^−/−^* and *5a^−/−^;5b^−/−^*larvae compared to wild-type siblings (WTs), indicating active mRNA degradation of *5a^−/−^* transcripts (Fig. S1C). Similarly, qPCR analysis showed that *5b* transcript levels were significantly reduced in *5b^−/−^* and *5a^−/−^;5b^−/−^*larvae compared to WTs at 5 dpf, also showing active mRNA degradation of *5b^−/−^*transcripts (Fig. S1D).

No significant gross morphological differences were observed between *slc13a5* mutants and WTs at 5 dpf (Fig. 1I-L). To examine neural changes, we performed immunostaining for the neuronal marker HuC/D and quantified the size of all three brain regions at 5 dpf. *slc13a5^−/−^* larvae exhibited 5.5% (*5a^−/−^*), 4.9% (*5b^−/−^*) and 6.3% (*5a^−/−^;5b^−/−^*) reduction in the MB size compared to WTs, with the forebrain and hindbrain size unchanged (Fig. 1M-P). Consistently, the effect of *slc13a5* mutations on MB size correlates with the MB expression of both paralogs at a later zebrafish developmental stage. Next, we performed a survival curve analysis (0-30 dpf) and observed a drastic mortality of *slc13a5^−/−^* larvae, with only 15% (*5a^−/−^*), 10.7% (*5b^−/−^*) and 5.4% (*5a^−/−^;5b^−/−^*) mutants surviving until 30 dpf and into adulthood (Fig. 1Q).

### *slc13a5^−/−^* larvae exhibit behavioral deficits, reduced neuronal numbers, and increased neuronal apoptosis

Individuals suffering from SLC13A5 epilepsy exhibit comorbidities including cognitive impairments and sleep challenges (12, 24). To determine the behavioral effect of *slc13a5* loss-of-function on zebrafish behavior, we performed a 10-minute acoustic startle protocol by exposing 5 dpf larvae to 440 Hz vibration pulses in the light and calculated the total distance travelled (Fig. 2A). The *slc13a5^−/−^* larvae moved significantly longer distances compared to WTs (413.2% *5a^−/−^*; 1213.3% *5b^−/−^*; 285.9% *5a^−/−^;5b^−/−^*), exhibiting hypersensitivity to acoustic startle and a lack of short-term habituation, which can be a measure of learning potential and might resemble the cognitive dysfunction in SLC13A5 individuals (Fig. 2B and C). The locomotion heatmaps further illustrate a hyperactivity phenotype of *slc13a5^−/−^*larvae compared to WTs when exposed to acoustic startle (Fig. 2D). Next, we assessed the sleep behavior of 5 dpf *slc13a5^−/−^*larvae by tracking the total distance moved during 10 hours of darkness in the night. The *slc13a5^−/−^*larvae traveled longer distances compared to WTs (163.4% *5a^−/−^*; 234.8% *5b^−/−^*; 148.4% *5a^−/−^;5b^−/−^*), showing disruptions in nighttime behaviors (Fig. 2E). We further examined swimming activity at 5 dpf for 30 minutes in the light and *slc13a5^−/−^* larvae travelled significantly longer distances compared to WTs (145.7% *5a^−/−^*; 119.2% *5b^−/−^*; 120% *5a^−/−^;5b^−/−^*), indicating a hyperactive phenotype during daytime (Fig. S2A), which is consistent with other epileptic zebrafish models (25). Approximately 20% of individuals with epilepsy suffer from anxiety (26, 27), thus we tested the “wall-hugging” thigmotaxis behavior of our zebrafish mutants, a validated measure of anxiety in animals and humans (28–30). We exposed the 5 dpf larvae to 6 minutes of light and 4 minutes of darkness, and measured the distance they travelled and time they spent close to the wall, *i.e*., in the outer well zone (Fig. S2B). There were no significant changes in the distance travelled and time spent close to the wall in mutants compared to WTs (Fig. S2C and D). Some individuals with epilepsy are more sensitive to stimuli causing reflex seizures (31), we thus challenged the 5 dpf larvae with an acute treatment of pentylenetetrazol (PTZ) to induce instantaneous convulsions, followed by exposure to thigmotaxis protocol as described above. Interestingly, the *slc13a5^−/−^* larvae swam significantly longer distances (101.2% *5a^−/−^*; 75.8% *5b^−/−^*; 85% *5a^−/−^;5b^−/−^*) and spent significantly more time hugging the wall (110% *5a^−/−^*; 63.6% *5b^−/−^*; 70.3% *5a^−/−^;5b^−/−^*) compared to WTs, indicating anxiety-like behavior in the presence of a chemical stimulus (Fig. S2E and F). Combined, *slc13a5^−/−^* larvae display a range of behavioral deficiencies.

**Fig. 2.**
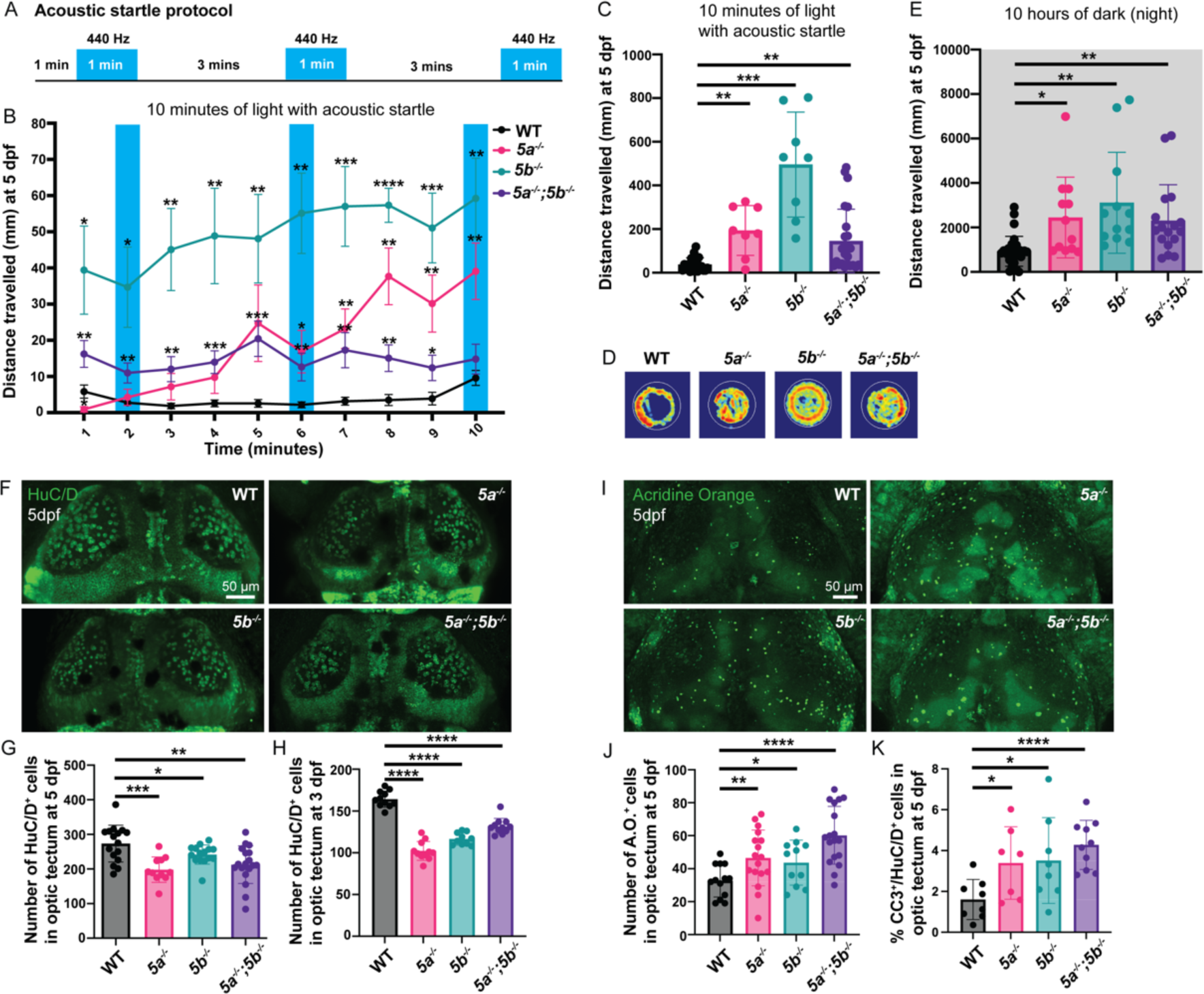
Analysis of startle response and sleep disturbances in *slc13a5* mutants, along with their neuron population and neuronal apoptosis assessment. (A) Schematic representation of acoustic startle protocol. (B, C) Quantification of distance travelled every one minute (B) and in 10 minutes (C) in acoustic startle at 5 dpf. *slc13a5* mutants are more responsive to startle with significantly higher distance travelled compared to WT. WT, n=21; *5a^−/−^,* n=8; *5b^−/−^*, n=8; *5a^−/−^;5b^−/−^*, n=24. (D) Heat maps of locomotion of larvae at 3 dpf under acoustic startle, showing that *slc13a5* mutants are more active than WTs (red shows highest presence and blue shows least presence). WT, n=4; *5a^−/−^,* n=4; *5b^−/−^*, n=4; *5a^−/−^;5b^−/−^*, n=4. (E) Quantification of distance travelled in 10 hours of darkness in night at 5 dpf. *slc13a5* mutants are more active than WTs, showing sleeping difficulties. WT, n=30; *5a^−/−^,* n=12; *5b^−/−^*, n=12; *5a^−/−^;5b^−/−^*, n=17. (F) 5 dpf WTs and *slc13a5* mutants; α-HuC/D (green). (G) Quantification of HuC/D^+^ cells in the optic tectum at 5 dpf. *slc13a5* mutants show a reduction in neuron numbers compared to WTs. WT, n=16; *5a^−/−^,* n=11; *5b^−/−^*, n=15; *5a^−/−^;5b^−/−^*, n=18. (H) Quantification of HuC/D^+^ cells in the optic tectum at 3 dpf. *slc13a5* mutants show a reduction in neuron numbers compared to WTs. WT, n=11; *5a^−/−^,* n=11; *5b^−/−^*, n=10; *5a^−/−^;5b^−/−^*, n=11. (I) Live 5 dpf *slc13a5* mutants and WTs, stained with A.O. (green). (J) Quantification of A.O.^+^ cells in the optic tectum at 5 dpf. *slc13a5* mutants show an increase in A.O.^+^ cell numbers compared to WTs. WT, n=13; *5a^−/−^,* n=17; *5b^−/−^*, n=11; *5a^−/−^;5b^−/−^*, n=18. (K) Percentage of CC3^+^/HuC/D^+^ cells (out of total HuC/D^+^ cells) in the optic tectum at 5 dpf. *slc13a5* mutants show an increase in CC3^+^ neuron population compared to WTs. WT, n=8; *5a^−/−^,* n=7; *5b^−/−^*, n=8; *5a^−/−^;5b^−/−^*, n=10. Data are Mean ± S.D., *P ≤ 0.05, **P ≤ 0.01, ***P ≤ 0.001, ****P ≤ 0.0001-Unpaired t-test. A.O., acridine orange; CC3, cleaved-Caspase3.

We further studied whether these *slc13a5^−/−^* larval behavioral deficits correlated with any molecular neural changes. We focused our assessments on the optic tectum region of the MB as its size was significantly affected in our mutants. We performed immunostaining for HuC/D to quantify the number of neurons and observed a reduction in 5 dpf *slc13a5^−/−^* larvae compared to WTs (−27.5%, *5a^−/−^*; −11.6%, *5b^−/−^*; −22.3%, *5a^−/−^;5b^−/−^*; Fig. 2F and G). To determine when the neuronal population started to decline, we quantified neuronal numbers (HuC/D positive) at 3 dpf and still observed a decrease in *slc13a5* mutants at this earlier stage (−37.7%, *5a^−/−^*; −29.1%, *5b^−/−^*; −19.9%, *5a^−/−^;5b^−/−^*; Fig. S3A, Fig. 2H), causing us to question if fewer neurons were born in *slc13a5^−/−^* to start. We quantified the number of neurons (HuC/D positive) around peak neurogenesis (28 hpf) in the region that gives rise to optic tectum and found no significant differences between *slc13a5^−/−^* larvae and WTs at this stage (Fig. S3B and C), leading us to question if cell death was a potential cause of the decreased neuronal numbers. We performed acridine orange (A.O.) staining at 5 dpf to label apoptotic cells and showed that *slc13a5^−/−^*larvae exhibited an increase in cell death (42.3% *5a^−/−^*; 33.5% *5b^−/−^*; 84.2% *5a^−/−^;5b^−/−^*) compared to WTs (Fig. 2I and J). This result was further validated by performing co-immunostaining for HuC/D and cleaved-Caspase 3 (CC3), a marker for apoptotic cells, at 5 dpf where we observed an increase in the percentage of CC3+ neurons in *slc13a5^−/−^* larvae compared to WTs (111.9%, *5a^−/−^*; 120%, *5b^−/−^*; 167.5%, *5a^−/−^; 5b^−/−^*), signifying that the reduction in neuronal numbers is a consequence of increased cell death (Fig. S3D and Fig. 2K).

### *slc13a5^−/−^* larvae experience excitation/inhibition imbalance, strong brain hyperexcitability, and compromised mitochondrial function

Next, in *slc13a5* mutant zebrafish we tested for disruption of neuronal excitatory/inhibitory (E/I) balance and synchronous hyperexcitation as found in mammalian (32, 33) and zebrafish (23, 34, 35) epilepsies. Using qPCR at 5 dpf, we quantified excitatory glutamatergic (*vglut2a*) and inhibitory GABAergic (*gad1b*) neuronal marker expression in larval brains and showed that *vglut2a* transcripts were significantly upregulated, whereas *gad1b* transcripts were significantly downregulated in *slc13a5^−/−^* larvae compared to WTs, suggestive of an E/I imbalance (Fig. 3A and B). Our group and others have shown previously that seizure induction in epilepsy animal models leads to an upregulated expression of immediate early genes (IEG) in the brain (20, 23, 36). We used qPCR at 5 dpf to quantify the expression of an IEG, *fosab,* and observed a significant upregulation in *fosab* transcripts in the *slc13a5^−/−^*larvae compared to WTs, further suggestive of brain hyperexcitability (Fig. 3C).

**Fig. 3.**
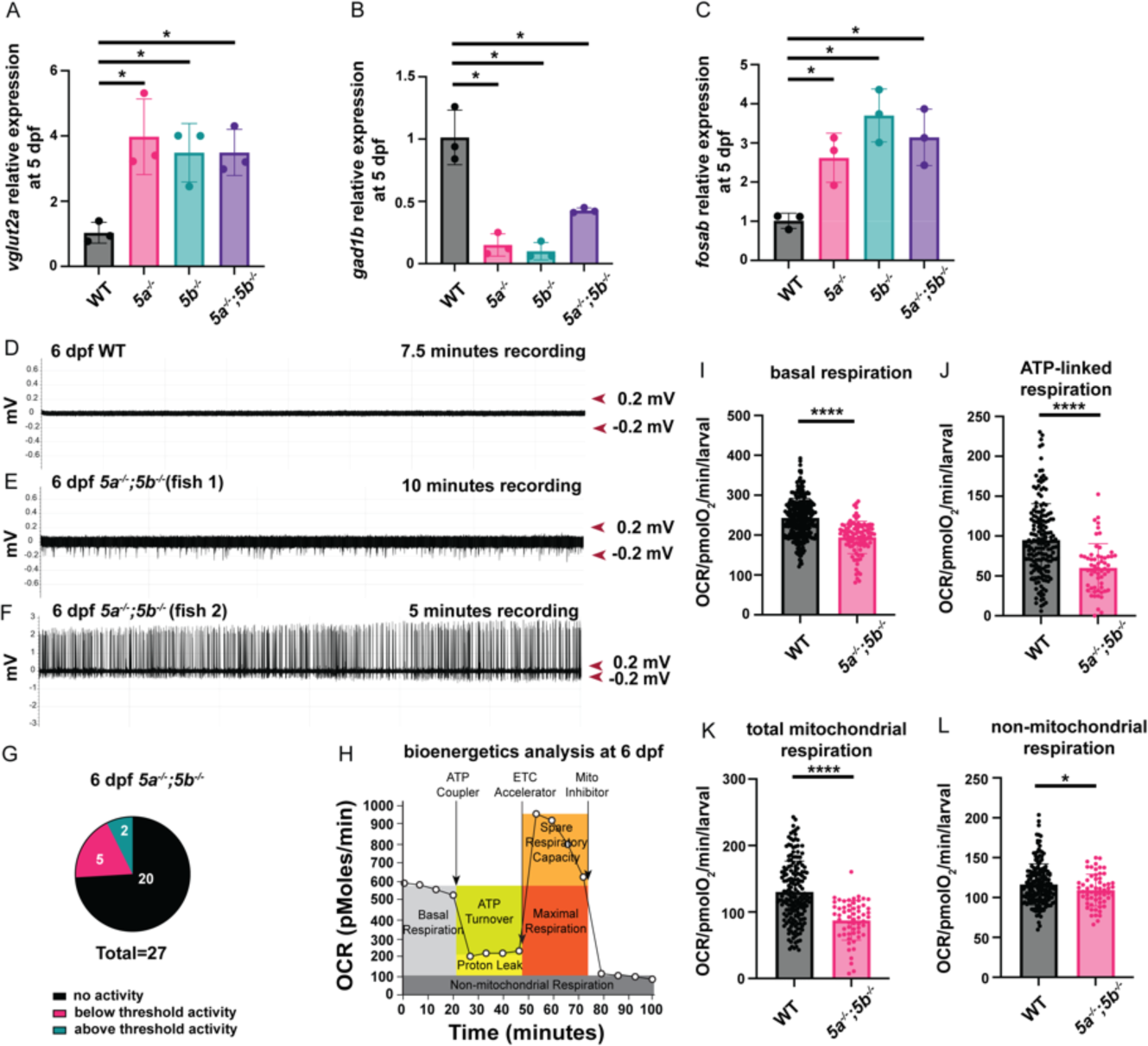
E/I imbalance, brain hyperexcitability and metabolic health analysis in *slc13a5* mutants. (A-C) qPCR analysis for relative *vglut2a, gad1b* and *fosab* mRNA expression in the heads of 5 dpf *slc13a5* mutants compared to WT. WT and *slc13a5* mutants, n = 3 × 10 larvae assessed as three biological and two technical replicates each. *vglut2a* is upregulated and *gad1b* is downregulated in *slc13a5* mutants indicating dysfunctional E/I balance. *fosab* is upregulated in *slc13a5* mutants indicating brain hyperexcitability. (D-G) Representative extracellular recordings obtained from optic tectum of 6 dpf WTs and *slc13a5* mutants, and a pie chart of number of *slc13a5* mutants showing different patterns of activity. The repetitive inter-ictal like discharges (<1s duration) with above threshold (>0.2mV), high-frequency, large-amplitude spikes seen in *slc13a5* mutants are indicative of increased network hyperexcitability. *slc13a5^−/−^* larvae also displayed below threshold (<0.2mV) brain activity. WT, n=8 out of 8 with no abnormal activity. (H) Schematic representation of how the Seahorse bioanalyser displays mitochondria bioenergetics being regulated by pharmacological inhibitors. (I) Quantification of basal respiration at 6 dpf. *slc13a5* mutants exhibit a significant reduction in basal respiration compared to WT. WT, n=58; *slc13a5* mutants, n=20 (individual values plotted from five cycles). (J) Quantification of ATP-linked respiration at 6 dpf. *slc13a5* mutants exhibit a significant reduction in ATP-linked respiration compared to WT. WT, n=59; *slc13a5* mutants, n=20 (individual values plotted from three cycles). (K) Quantification of total mitochondrial respiration at 6 dpf. *slc13a5* mutants exhibit a significant reduction in total mitochondrial respiration compared to WT. WT, n=59; *slc13a5* mutants, n=20 (individual values plotted from three cycles). (L) Quantification of non-mitochondrial respiration at 6 dpf. *slc13a5* mutants exhibit a significant reduction in non-mitochondrial respiration compared to WT. WT, n=59; *slc13a5* mutants, n=20 (individual values plotted from three cycles). Data are Mean ± S.D., *P ≤ 0.05, ****P ≤ 0.0001-Unpaired t-test.

Next, to characterize seizure-like events, we measured extracellular field potentials from the optic tectum of agarose-immobilized 6 dpf WT and *slc13a5^−/−^* larval brains. WTs showed no evidence of abnormal brain activity (Fig. 3D); however, extracellular field recordings in 30% (*5a^−/−^*), 10% (*5b^−/−^*) and 7.4% (*5a^−/−^;5b^−/−^*) of *slc13a5^−/−^* larval brains revealed repetitive inter-ictal like discharges (<1s duration) with high-frequency, large-amplitude, above threshold (>0.2mV) spikes, consistent with a spontaneous epileptic phenotype (Fig. S4A-C, Fig. 3E-G). Notably, 40% (*5a^−/−^*), 50% (*5b^−/−^*) and 18.5% (*5a^−/−^;5b^−/−^*) of *slc13a5^−/−^* larvae also displayed repetitive but below threshold (<0.2mV) brain activity (Fig. 3G, Fig. S4C). No longer duration (>1s) discharges characterized as ictal-like events (37) were identified in the mutants. We speculate that a range of electrical activities, i.e., above and below threshold spikes, observed in our mutants could be because of the stochastic nature of seizures (38). Moreover, electrophysiological recordings provide only a snapshot of the brain’s electrical activity at the location and time of recording, leading to some spatio-temporal restrictions of this approach.

Dysregulation of mitochondrial function is observed across various types of epilepsy, including in zebrafish models (23, 39-41) and here we used the Seahorse XF Flux Bioanalyzer to assess bioenergetics in our *slc13a5^−/−^*larvae at 6 dpf (Fig. 3H). We observed a dampening in basal respiration (−20.1%), ATP-linked respiration (−36.6%), total mitochondrial respiration (−32.9%), and non-mitochondrial respiration (−6.1%) in *5a^−/−^;5b^−/−^* larvae compared to WTs (Fig. 3I-L). In contrast, maximum respiratory capacity and reserve capacity were upregulated by 28.2% and 351.2%, respectively (*5a^−/−^;5b^−/−^*), whereas proton leaks were unchanged (Fig. S5A-C), consistent with other reports whereby the loss of citrate uptake by cells due to *SLC13A5* mutations causes changes in cellular metabolism (42–45).

### Knockdown of NMDA receptor subunit rescues dysregulated calcium activity, neurometabolism, and startle behaviors in *slc13a5^−/−^* larvae

The underlying mechanism linking citrate transporter mutations to seizures remains debated (11), with one hypothesis that the accumulation of extracellular citrate causes chelation of ions that results in E/I imbalance. N-methyl-D-aspartate (NMDA) receptors are glutamate-gated channels that allow the timely influx of calcium (Ca^+2^) to mediate excitatory signaling. Zinc is a critical NMDA receptor inhibitor that helps regulate Ca^+2^ uptake and prevent prolonged excitation. Citrate is a potent zinc chelator, which when present in excess might bind all the zinc and prevent it from modulating NMDA receptor function, causing unremitting Ca^+2^ influx and excitatory signaling.

First, we tested whether *slc13a5* mutants display increased Ca^+2^ events, as would be predicted if NMDA receptor signaling was overactive. We crossed WT and *5a^−/−^;5b^−/−^* larvae with the *Tg(elavl3:Hsa.H2B-GCaMP6s)* zebrafish that express a neuron-specific calcium biosensor, and performed live calcium imaging at 3 dpf. *5a^−/−^;5b^−/−^* larvae displayed an increase of 19% in the amplitude and 469% in the frequency of calcium events compared to WTs (Fig. 4A and B, Movies S1 and S2). The representative single neuron calcium traces illustrate the extent of increased Ca^+2^ signaling in the *5a^−/−^;5b^−/−^* mutants (Fig. 4C). Next, we assessed phosphorylated ERK (pERK) levels, an indirect downstream reporter of calcium signaling. We co-stained with pERK- and total ERK-(tERK) selective antibodies and imaged 5 dpf larval brains. Using a standard zebrafish reference brain to calculate normalized pERK levels as described previously (46, 47), whole-brain activity maps were generated with voxels exhibiting significantly higher (green) and lower (magenta) pERK levels in the different brain regions. We found a significant increase in pERK signaling (green) in the forebrain, MB, and hindbrain in *5a^−/−^;5b^−/−^* mutants compared to WTs, signifying upregulation of ERK phosphorylation possibly due to enhanced calcium influx in the neurons (Fig. 4D). Notably, the lower pERK intensity observed in the eyes (magenta) could be due to autofluorescence while imaging and therefore we do not consider it further.

**Fig. 4.**
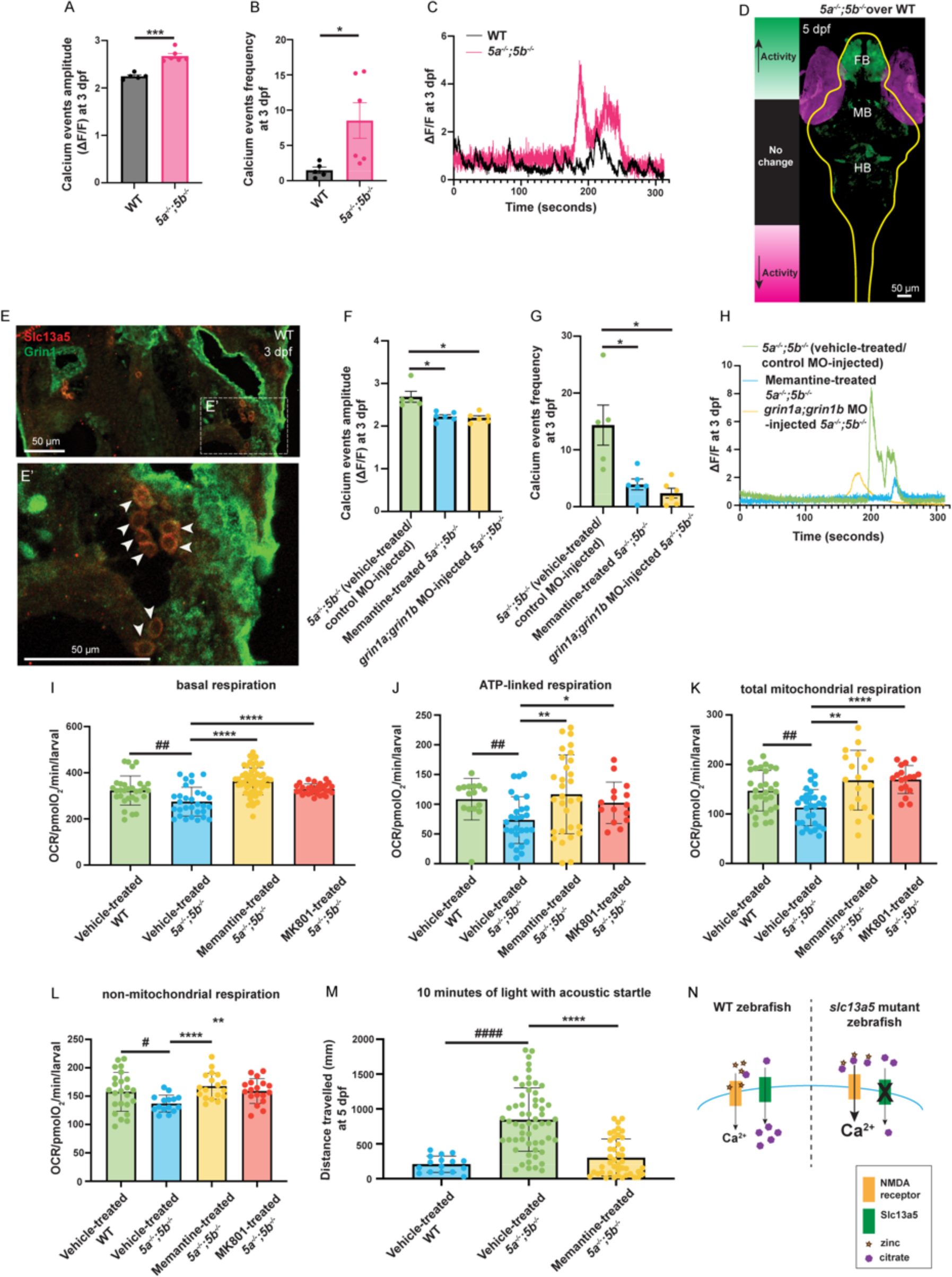
Calcium event analysis and assessment of the effect of suppressing NMDA receptor signaling in *slc13a5* mutants. (A-C) Quantification of the amplitude and frequency of calcium events (ΔF/F>2) and representative single neuron calcium traces at 3 dpf. *5a^−/−^;5b^−/−^* larvae show significant increase in calcium events compared to WTs. WT, n=5; *5a^−/−^;5b^−/−^,* n=6. (D) Whole-brain activity MAP-map depicting significant changes in pERK signal, calculated using Mann-Whitney U statistic Z score. The significance threshold was set based on a FDR where 0.05% of control pixels would be called as significant. Voxels exhibiting significantly higher intensity values of pERK are denoted in green and those exhibiting significantly lower pERK intensity values are depicted in magenta, in the *slc13a5* mutants compared to WTs at 5 dpf. *slc13a5* mutants show an enhanced neural activity via increased pERK levels in different regions of the brain (green) compared to WT. The lower pERK intensity shown in the eyes (magenta) could be a background noise captured while imaging. WT, n=9; *5a^−/−^;5b^−/−^*, n=8. (E) Section of 3 dpf MB region of WTs; α-Slc13a5, α-Grin1. (E’) Higher magnification of dashed box in E. White arrowheads point to Slc13a5^+^/Grin1^+^ cells in MB. (F-H) Quantification of the amplitude and frequency of calcium events (ΔF/F>2) and representative single neuron calcium traces at 3 dpf. Memantine treatment and *grin1a;grin1b* MO injections significantly reduced the amplitude and frequency of calcium events in *5a^−/−^*;*5b^−/−^* larvae compared to vehicle-treated or control MO-injected *5a^−/−^*;*5b^−/−^* larvae. Vehicle-treated/control MO-injected *5a^−/−^*;*5b^−/−^*larvae, n=5; memantine-treated *5a^−/−^*;*5b^−/−^* larvae, n=6; *grin1a;grin1b* MO-injected *5a^−/−^*;*5b^−/−^*larvae, n=6. (I) Quantification of basal respiration before and after treatment with memantine and MK801 at 6 dpf. Memantine and MK801 treatments rescue compromised basal respiration in *5a^−/−^;5b^−/−^* larvae. Vehicle-treated WT, n=6; vehicle-treated *5a^−/−^;5b^−/−^*, n=6; memantine-treated *5a^−/−^;5b^−/−^*, n=12; MK801-treated *5a^−/−^;5b^−/−^*, n=6 (individual values plotted from five cycles). (J) Quantification of ATP-linked respiration before and after treatment with memantine and MK801 at 6 dpf. Memantine and MK801 treatments rescue compromised ATP-linked respiration in *5a^−/−^;5b^−/−^* larvae. Vehicle-treated WT, n=5; vehicle-treated *5a^−/−^;5b^−/−^*, n=9; memantine-treated *5a^−/−^;5b^−/−^*, n=10; MK801-treated *5a^−/−^;5b^−/−^*, n=5 (individual values plotted from three cycles). (K) Quantification of total mitochondrial respiration before and after treatment with memantine and MK801 at 6 dpf. Memantine and MK801 treatments rescue compromised total mitochondrial respiration in *5a^−/−^;5b^−/−^* larvae. Vehicle-treated WT, n=10; vehicle-treated *5a^−/−^;5b^−/−^*, n=10; memantine-treated *5a^−/−^;5b^−/−^*, n=6; MK801-treated *5a^−/−^;5b^−/−^*, n=6 (individual values plotted from three cycles). (L) Quantification of non-mitochondrial respiration before and after treatment with memantine and MK801 at 6 dpf. Memantine and MK801 treatments rescue compromised non-mitochondrial respiration in *5a^−/−^;5b^−/−^*larvae. Vehicle-treated WT, n=8; vehicle-treated *5a^−/−^;5b^−/−^*, n=5; memantine-treated *5a^−/−^;5b^−/−^*, n=6; MK801-treated *5a^−/−^;5b^−/−^*, n=6 (individual values plotted from three cycles). (M) Quantification of total distance travelled in 10 minutes in acoustic startle before and after treatment with memantine at 5 dpf. Memantine treatment rescues impaired startle response in *5a^−/−^;5b^−/−^* larvae. Vehicle-treated WT, n=16; vehicle-treated *5a^−/−^;5b^−/−^*, n=58; memantine-treated *5a^−/−^;5b^−/−^*, n=43. (N) Model of relationship between *slc13a5* mutations and NMDA receptor signaling: Unlike in WT larvae, *slc13a5* mutations in zebrafish presumably cause an increase in extracellular citrate levels that lead to zinc chelation, thereby preventing the inhibitory effect of zinc on NMDA receptors. This ultimately leads to an excessive influx of calcium inside the neurons, a plausible cause of epileptic phenotypes in the *slc13a5* mutants. Data are Mean ± S.E.M. and Mean ± S.D., *P ≤ 0.05, ^#^P ≤ 0.05, **P ≤ 0.01, ^##^P ≤ 0.01, ***P ≤ 0.001, ****P ≤ 0.0001, ^####^P ≤ 0.0001-Unpaired t-test. MO, morpholino. FDR, false discovery rate.

Next, we investigated the involvement of NMDA receptor signaling in *5a^−/−^;5b^−/−^* larvae phenotypes using pharmacological and genetic approaches. Pharmacologically, we exposed WT and mutant zebrafish larvae to memantine, an established NMDA antagonist used in other zebrafish studies (48). Genetically, we designed morpholinos to knockdown the NMDA receptor subunit *grin1* (also known as *glun1*), reported to be responsible for calcium permeability of NMDA receptors (49, 50). Both *grin1a* and *grin1b* paralogs are expressed in the zebrafish brain and their loss of function affects behaviors (51, 52). We confirmed Grin1 expression in Slc13a5^+^ MB neurons by immunostaining 3 dpf larvae and observed their co-expression in several cells in the MB (Fig. 4E, E’). We co-injected *grin1a;grin1b* morpholinos (MOs) in the WT and *5a^−/−^;5b^−/−^* zebrafish, expressing the *Tg(elavl3:Hsa.H2B-GCaMP6s)* transgene and performed live calcium imaging at 3 dpf. 50 μM memantine-treatment for 2 hours prior to imaging and *grin1a;grin1b* MO injection in *5a^−/−^;5b^−/−^*larvae caused a reduction of −17.1% (memantine-treated) and −18.2% (*grin1a;grin1b* MO injected) in the amplitude and −72.7% (memantine-treated) and −83.3% (*grin1a;grin1b* MO injected) in the frequency of calcium events compared to vehicle-treated (0.5% DMSO) or control MO-injected *5a^−/−^;5b^−/−^* larvae (Fig. 4F and G, Movies S3-S5). The representative single neuron calcium traces illustrated that *5a^−/−^;5b^−/−^* larvae exhibited fewer above-threshold events after memantine-treatment or *grin1a;grin1b* MO injection (Fig. 4H). Notably, there were no significant changes in the amplitude or frequency of calcium events in WT larvae after treatment with memantine or *grin1a;grin1b* MO injections (Fig. S6A and B). Importantly, a significant reduction in the Grin1 fluorescence intensity, as measured by calculating corrected total cell fluorescence (CTCF) in the brains of Grin1 immunostained *grin1a;grin1b* MO-injected WTs compared to control MO-injected WTs at 3 dpf, was observed, confirming efficacy of *grin1a;grin1b* MOs (Fig. S6C). These results demonstrate a role for NMDA receptors in *slc13a5* mutant dysregulation of Ca^+2^ signaling.

An alternative mechanistic hypothesis is that the loss of *intracellular* citrate disrupts neuronal homeostasis, leading to hyperexcitability. If this alternative hypothesis is true, then blocking NMDA receptors should have no additional effect in the mutants on a key measure of neuronal homeostasis: bioenergetics. 6 dpf *5a^−/−^;5b^−/−^* larvae were treated with 50 μM memantine, 50 μM MK801, also a potent NMDA antagonist (52) or vehicle (0.5% DMSO) for 2 hours prior to initiating the Seahorse assay (as per above). Blocking NMDA receptors significantly rescued several bioenergetics parameters dysregulated in the *5a^−/−^;5b^−/−^* larvae, including basal respiration, ATP-linked respiration, total mitochondrial respiration, and non-mitochondrial respiration (Fig. 4I-L), suggesting that an interplay between extracellular citrate and NMDA receptors mediates the seizure-like phenotype in *slc13a5* mutants. Notably, these antagonists had no effect on these metabolic parameters in the WTs (Fig. S7A-D).

Finally, to show that NMDA receptors function affects circuit level behaviors of *slc13a5* mutant zebrafish, we examined whether the impaired startle behavior of our mutants was rescued by NMDA antagonism. We treated 5 dpf larvae with memantine (50 μM) or vehicle (0.5% DMSO) for 2 hours, followed by the acoustic startle protocol (as per above). Treatment with memantine rescued the impaired startle response in *5a^−/−^;5b^−/−^* larvae, causing a significant reduction in the total distance swam during the complete protocol (Fig. 4M). No effect was observed of memantine exposure on the swimming activity of the WTs in this acoustic startle protocol (Fig. S7E).

Taken together, we propose a model whereby deficiencies in Slc13a5 cause accumulation of extracellular citrate that excessively chelates zinc, which fail to efficiently buffer NMDA receptor activity leading to increased Ca^+2^ uptake and excessive excitatory firing that manifests as seizures (Fig. 4N).

## Discussion

It remains unclear why children with mutations in *SLC13A5* develop severe epilepsy and associated comorbidities. In the present study we created genetic loss-of-function mutants of both the paralogs of human *SLC13A5*, i.e., *5a* and *5b* in the zebrafish to facilitate the study of the underlying disease etiology. These *slc13a5* zebrafish mutants recapitulate many phenotypes observed in *SLC13A5*-deficient individuals, making this zebrafish model a robust and reliable tool. Mechanistically, we show that pharmacological inhibition or genetic knockdown of NMDA receptor signaling rescues the epileptic phenotypes, serving as the first empirical evidence for the ‘chelation hypothesis’ of SLC13A5 epilepsy. Combined, these zebrafish mutants are a translatable model to study underlying disease biology and may open new avenues of research towards novel treatment options for this disorder.

We first characterized the *slc13a5* mutants for disruption in various behaviors, which is a common strategy to test for gross neural network defects. Deficiencies in behaviors were observed across several assays in *slc13a5* mutants, including those that recapitulate the sleep disturbances and cognitive disabilities (by showing impaired learning of the startle response) observed in *SLC13A5* individuals, further demonstrating the applicability of this model system. To investigate whether these behavioral phenotypes correlate with molecular changes in mutant brains, we focused our characterization to the optic tectum given the high level of *slc13a5* expression in the MB. There was a significant reduction in the number of neurons and MB size, along with a concomitant increase in the neuronal death in our *slc13a5* mutants. Microcephaly is reported in ∼10% of SLC13A5 individuals (12, 24), indicating that brain size reduction (a possible consequence of cell death) is present but is less penetrant in humans. We speculate that this increased apoptosis is a consequence of seizure-like events in the mutant brain, since other animal model and patient data show that epileptic seizures cause neuronal death via activation of apoptosis or autophagy signaling pathways (34, 53-55). It is also possible that a block in citrate transport causes a loss of intracellular citrate, a crucial metabolite for energy production, and thus alters neuronal metabolic health (43). Our zebrafish *slc13a5* mutants exhibited compromised neurometabolism, consistent with a metabolic phenotype. And finally, it is notable that only a small percentage of the zebrafish mutants survive to adulthood, suggesting that perhaps disease progression becomes incompatible with survival in the zebrafish.

Brain MRI and EEG recordings from SLC13A5 epilepsy patients show punctate white matter lesions and abnormal brain activity, respectively (56, 57). Here, the electrical field recordings, as well as the upregulation of the immediate early gene, *fosab,* and perturbed E/I balance, is consistent with an epilepsy phenotype in our *slc13a5* mutants. This demonstration of seizure-like activity is not trivial given that citrate transporter mutations in *C. elegans, Drosophila,* and mouse fail to robustly produce epileptiform events. That said, nearly three-fourths of the zebrafish *5a^−/−^;5b^−/−^* mutants displayed no hyperexcitability when assessed by extracellular field recordings, which is unexpected. This percentage of animals displaying no activity is considerably smaller in single *5a^−/−^* or *5b^−/−^* mutants – one-quarter and one third, respectively, raising questions as to why the double mutant is neuroprotective. Furthermore, this finding is unexpected given that across all other measures the double mutants displayed strong phenotypes, including behavioral assays, molecular analysis, bioenergetics profiling and especially Ca^+2^ flux that also is a measure of neuronal activity. Perhaps the optic tectum is not the focal point for the generation of these seizure-like events in the double mutants, but further studies are needed.

A key finding of our studies is the mechanistic evidence suggesting an interplay between a defective citrate transporter and activation of NMDA receptors. Little is known about the molecular etiology of this disorder and one of the hypotheses is that excessive extracellular citrate levels due to the loss of transmembrane transport chelates free zinc and prevents its binding to NMDA receptor subunits, thereby causing an uncontrolled influx of calcium into the neurons and leading to increased seizure susceptibility in the patients (11). We show that the *slc13a5* mutants exhibit upregulated levels of pERK, a downstream reporter of calcium activity, as well as a significant increase in calcium events as revealed by calcium imaging in live larvae. Interestingly, both memantine treatment as well as *grin1a;grin1b* MO injections led to a significant normalization of the amplitude and frequency of calcium events in the *slc13a5* mutants, demonstrating a role for NMDA receptors in *slc13a5* hyperexcitable phenotypes. We also show that pharmacological inhibition of NMDA signaling using memantine and MK801 improves the metabolic profiles of our mutants, further validating a possible relationship between extracellular citrate and NMDA receptors. Further, memantine treatment rescued the impaired acoustic startle response of our mutant zebrafish, demonstrating a functional interaction can rescue animal behaviors as well. Combined, these findings suggest that inhibition of NMDA receptor signaling could be a potential target for the treatment of epileptic seizures in humans.

A potential limitation of these studies is the expression of Slc13a5 in the plasma membrane of both neurons and astrocytes in zebrafish. In the human brain, SLC13A5 is expressed primarily in neurons (1, 2, 11), whereas in mice it is expressed in both neurons and astrocytes (58–60), which was considered a confounding factor when translating back to humans. And now, in zebrafish we show that Slc13a5 also is expressed in both neurons and astrocytes, perhaps reflecting a functional difference in metabolic requirements for citrate by neural cells across vertebrate species. To circumvent any issues related to the dual expression of this citrate transporter in SLC13A5 epilepsy and to overcome any potential shortcomings of using zebrafish in isolation (61), induced pluripotent stem cells can be established from SLC13A5 individuals and used to generate human brain organoids. These brain organoids can be employed to validate these zebrafish findings in a human model and to overexpress *SLC13A5* to further shed insights into the function of SLC13A5 in the brain. In fact, two recent studies show that *Slc13a5* overexpression causes autism and progeria-like phenotypes in mice models (62, 63).

In conclusion, *slc13a5* mutant zebrafish are a promising model for the discovery of disease biology and a new tool for high-throughput drug screening and the testing of candidate drugs for this disorder.

## Materials and methods

### Zebrafish maintenance

Adult zebrafish (TL and AB strains) were maintained at 28°C in a 14-hour light/10-hour dark cycle under standard aquaculture conditions, and fertilized eggs were collected via natural spawning. The animals were fed twice daily with Artemia. Zebrafish embryos and larvae were maintained in embryo medium in a non-CO_2_ incubator (VWR) at 28°C on the same light–dark cycle as the aquatic facility. All protocols and procedures were approved by the Health Science Animal Care Committee (protocol number AC22-0153) at the University of Calgary in compliance with the Guidelines of the Canadian Council of Animal Care.

### Generation of zebrafish mutants by CRISPR/Cas9

*5a^−/−^* and *5b^−/−^* zebrafish were generated by using CRISPR/Cas9 mutagenesis. The identified founders carried 11-nucleotide deletion in fifth exon for *5a* mutations and one-nucleotide substitution followed by 13-nucleotide insertion in fifth exon for *5b* mutations. The gRNA sequences are CCACACTCACAGGCACTGGACCA for *5a* mutation and GTTTGCTATGCTGCCAGTGTTGG for *5b* mutation. WTs, single homozygous (*5a^−/−^* and *5b^−/−^*) and double homozygous (*5a^−/−^;5b^−/−^*) larvae were used for the experiments (by breeding heterozygous adults). Genotyping to distinguish *5a^−/−^, 5a^+/−^* and WTs was carried out by performing high resolution melt analysis (HRMA) using primers listed in Table S1. Genotyping to distinguish *5b^−/−^, 5b^+/−^* and WTs was carried out by performing HRMA using primers listed in Table S1, followed by running the non-heterozygous PCR products on agarose gel, based on different sizes of PCR product bands (WTs−118 bp and *5b^−/−^*−131 bp). Survival of animals was analyzed by recording their mortality rate every other day beginning from 0 dpf until 30 dpf and percentage survival was plotted (GraphPad Prism).

### Morpholino injections

The *grin1a* (5’-CCAGCAGAAGCAGACGCATCGT-3’) and *grin1b* (5’-CGAACAGAACCAGGCGCATTTTGCC-3’) MOs were purchased from Genetools (Philomath, OR) and co-injected at the one-cell stage at 0.75 ng concentrations in all experiments described. The standard control MO (5’-CCTCTTACCTCAGTTACAATTTATA-3’) purchased from Genetools was also injected at 0.75 ng concentration. The doses of all MOs were determined as optimal by titration (no toxic effects were observed).

### Gene expression analysis by in situ hybridization and qPCR

Digoxigenin-labeled probes were synthesized for *5a* and *5b* as described previously (64), using the primers listed in Table S2. The embryos and larvae were fixed at desired stages in 4% paraformaldehyde (Sigma-Aldrich) overnight at 4°C. The whole mount in situ hybridization protocol was performed as described previously (64). qPCR was performed for *5a*, *5b*, *vglut2a*, *gad1b,* and *fosab* on cDNA obtained from the heads of 5 dpf larvae. *rpl13* was used as internal control. The primers used for qPCR are listed in Table S3.

### Behavioral assays

Zebrafish larvae maintained in 48-well plates were habituated for 20 minutes, under ambient light. This was followed by behavioral assessment to measure distance moved in 100% light and during acoustic startle, using Zebrabox (Viewpoint Life Sciences). Similarly, larvae were habituated in 20 minutes of darkness, followed by tracking their movement in 100% darkness. For thigmotaxis protocol, we used 24-well plates and generated a protocol to divide each well into inner well zone and outer well zone. The larvae were habituated for 20 minutes in ambient light (or treated with 5 mM PTZ prior to this). This was followed by tracking the distance travelled and time spent close to the wall under 6 minutes of 100% light and 4 minutes of 100% darkness. Tracking of total distance moved and time taken as a measure of swimming behavior was analyzed using Zebralab V3 software (Viewpoint Life Sciences). Locomotion heatmaps depicting the movement of larvae were also generated by using Zebralab V3 software. All the behavioral assessments were performed on 5 dpf larvae.

### Drug treatments

Memantine (Cayman Chemical) and MK801 (Sigma-Aldrich) stock solutions were prepared in DMSO. The drugs were assessed for toxicity and the highest concentration which did not induce toxicity, was used in subsequent assays. On the day of experiments, the stock solutions were diluted to a final concentration of 50 μM in embryo medium. The zebrafish larvae were treated with 50 μM drugs for 2 hours, followed by behavior assessment, Seahorse assay and calcium imaging. The final DMSO concentration was 0.5% and was used as vehicle control in the drug treatment experiments.

### Electrophysiological measurements

Electrophysiological recordings in zebrafish larvae were performed as described previously (40, 65). Briefly, 6 dpf larvae were paralyzed using α-bungarotoxin (1 mg/ml, Tocris) and embedded in 1.2% low melting agarose (LMA). The dorsal side of the larvae was exposed to the agarose gel surface and accessible for electrode placement. Larvae were placed on an upright stage of an Olympus BX51WI upright microscope, visualized using a 5X MPlanFL N Olympus objective, and perfused with embryo media. A glass microelectrode (3–8 MΩ) filled with 2 M NaCl was placed into the optic tectum of zebrafish, and a recording was performed in current-clamp mode, low-pass filtered at 1 kHz, high-pass filtered at 0.1 Hz, using a digital gain of 10 (Multiclamp 700B amplifier, Digidata 1440A digitizer, Axon Instruments) and stored on a PC computer running pClamp software (Axon Instruments). Baseline recording was performed for ≥ five minutes. The threshold for detection of spontaneous events was set at three times noise, as described previously (66). The frequency and amplitude of spontaneous events were analyzed using the Clampit software 11.0.3 (Molecular Devices, Sunnyvale, CA).

### Metabolic measurements

Oxygen consumption rate (OCR) measurements were performed using the XF24 Extracellular Flux Analyzer (Seahorse Biosciences). Individual 6 dpf zebrafish larvae were placed in 24 wells on an islet microplate and an islet plate capture screen was placed over the measurement area to maintain the larvae in place. Seven measurements were taken to establish basal rates, followed by treatment injections and 18 additional cycles (67). Rates were determined as the mean of two measurements, each lasting 2 min, with 3 min between each measurement (68). Three independent assays were performed to establish metabolic measurements. A script was generated on R (using dplyr, reshape2, tidyr, purr and tidyverse libraries) to calculate the metabolic measurements for all the parameters.

### Immunofluorescence

To perform whole mount immunostaining (for HuC/D), 3 dpf and 5 dpf larvae were fixed in 4% paraformaldehyde overnight at 4°C, depigmented with a 1% H_2_O_2_/3% KOH solution, followed by antigen retrival with 150mM Tris HCl (pH 9.0) for 15 minutes at 68°C, permeabilization (PBS/0.3%Triton-X/1%DMSO/1%BSA/0.1%Tween-20) for 2 hours at room temperature (RT) and blocking (PBS/1%DMSO/2%fetal bovine serum (FBS)/1%BSA/0.1%Tween-20) for 1 hour at RT. Later, the larvae were incubated in primary antibody overnight at 4°C, followed by washing and secondary antibody incubation overnight at 4°C. Finally, the immunostained larvae were washed and counterstained with Hoechst (Thermo Fisher Scientific). pERK/tERK whole mount immunostaining on 5 dpf larvae were performed as described previously (47). The larvae were mounted in 0.8% LMA for imaging. To perform immunostaining (for HuC/D, Slc13a5, Gfap, cleaved-Caspase3 and Grin1) on 12 µm thick cryosections of 28 hpf embryos, 3 dpf and 5 dpf larvae, antigen retrival was carried out with citrate buffer (pH 6.0) for 15 minutes at 95°C. This was followed by permeabilization with 1% Triton-X for 30 minutes at RT and blocking in 5% normal donkey serum or 5% normal goat serum for 1 hour at RT. Later, the sections were incubated in primary antibody overnight at 4°C, followed by washing and secondary antibody incubation for 3 hours at RT. Finally, the immunostained sections were washed, counterstained with Hoechst and mounted for imaging. The primary antibodies (and their working concentrations) used in this study are listed in Table S4. Secondary antibodies were used at 1:200 concentration (Life Technologies).

### Live acridine orange staining

The larvae were raised in embryo medium containing 1-phenyly-2-thiourea (PTU) to prevent pigment formation. 5 dpf larvae were incubated for 30 minutes in embryo medium containing 0.002% A.O. solution (Sigma-Aldrich) at 28°C. The stained fish were washed with embryo medium several times and mounted in 0.8% LMA for imaging.

### Imaging and quantification

Bright-field images of in situ hybridization-stained embryos and larvae (whole mount and 12 µm thick cryosections) and live animals were obtained using Stereo Discovery.V8 microscope (ZEISS). The live A.O. stained/immunostained whole mount samples and cryosections were imaged at 20X magnification using LSM 880/LSM 900 confocal microscopes (ZEISS). 5 dpf larvae immunostained with pERK/tERK were imaged at 10X magnification in the Airyscan FAST mode of LSM 880. After imaging, the acquired confocal z-stacks were processed, cell counting and brain size measurements were performed with ZEN software. Fluorescence intensity measurements (for Grin1 signal) were performed by calculating CTCF using Fiji software.

### Brain activity analysis using pERK/tERK immunostaining

The processing and analysis pipeline is performed using the protocol described previously (46). Briefly, images were registered to a standard zebrafish reference brain using Computational Morphometry Toolkit (CMTK). The pERK images were normalized with the tERK staining. In MATLAB (MathWorks), the Mann-Whitney U statistic Z score is calculated for each voxel, comparing between the mutant and WT groups using MapMAPPING. The significance threshold was set based on a false discovery rate (FDR) where 0.05% of control pixels would be called as significant.

### Calcium imaging and data analysis

*Tg(elavl3:Hsa.H2B-GCaMP6s)^jf5Tg^* animals imported from Zebrafish International Resource Center were outcrossed with *5a^+/−^;5b^+/−^*to raise adults and subsequently *5a^−/−^;5b^−/−^* and WT larvae were obtained for calcium imaging. The larvae were raised in embryo medium containing PTU to prevent pigment formation. Briefly, 3 dpf larvae were immobilized using α-bungarotoxin and tricaine and embedded in 0.8% LMA. Larvae were imaged at 10X magnification the Airyscan FAST mode of LSM 880 and single-plane movies were acquired at 9.6 frames per second for ∼5 minutes. Mean grey value measurements of individual neurons in the MB were performed in Fiji and grey values were normalized against a basal mean grey value. A script was generated on R to calculate fluorescence variation measurements (ΔF/F) in each frame of all the neurons, followed by calculating the frequency and amplitude of calcium events (with a threshold value of ΔF/F>2) for each larvae recorded.

### Statistical analysis

GraphPad Prism software was used to perform statistical analysis. Data are represented as mean ± S.E.M. and mean ± S.D. p-values were calculated by unpaired t test.

### Author contributions

D.D. designed the study, performed the experiments, analyzed the data and wrote the manuscript; V.A.P. performed survival assay and some of the molecular assays; C.G. performed electrophysiological experiments; N.D. performed in situ hybridization and some behavioral/molecular assays; K.I. performed some bioenergetics experiments; D.M.K. designed the study, analyzed the data, edited the manuscript and supervised the work. All the authors commented on the manuscript.

## Acknowledgments

We would like to thank Boogyung Seo for help with behavioral analyses. We also thank Rachel Lacroix for training on pERK/tERK data processing. We further thank Sisu Han, Faizan Malik, Eric Samarut and Andrew Boyce for suggestions and training on calcium imaging processing. We also thank Pia Svendsen for training on confocal microscopes. We thank Ankita for generating Seahorse and calcium imaging data analysis scripts. We also thank Peng Huang lab for sharing their microinjection facility. We further thank Arthur Omorogiuwa and Natasha Klenin for assistance with zebrafish husbandry and reagent preparation, respectively. We thank Alicia Vandenbrink for administrative support. Finally, we thank TESS Research Foundation and their families for their continued interest of our zebrafish program.

## Funding sources

This project was funded by TESS Research Foundation Early-Career Investigator Research Grant to D.D. and by Brain Canada Platform Support Grant to D.M.K.

## Conflict of interest

D.M. Kurrasch is a co-founder of Path Therapeutics, an early-stage biotech focused on the development of novel drugs for the treatment of Dravet Syndrome. The work of D.M.K. with Path Therapeutics is not in conflict with the project presented herein. The other authors have nothing to disclose.

## Supplementary material

Supplementary material is available online.

## Supplementary Figures, Figure Legends, Movie Legends and Tables

**Figure S1.**
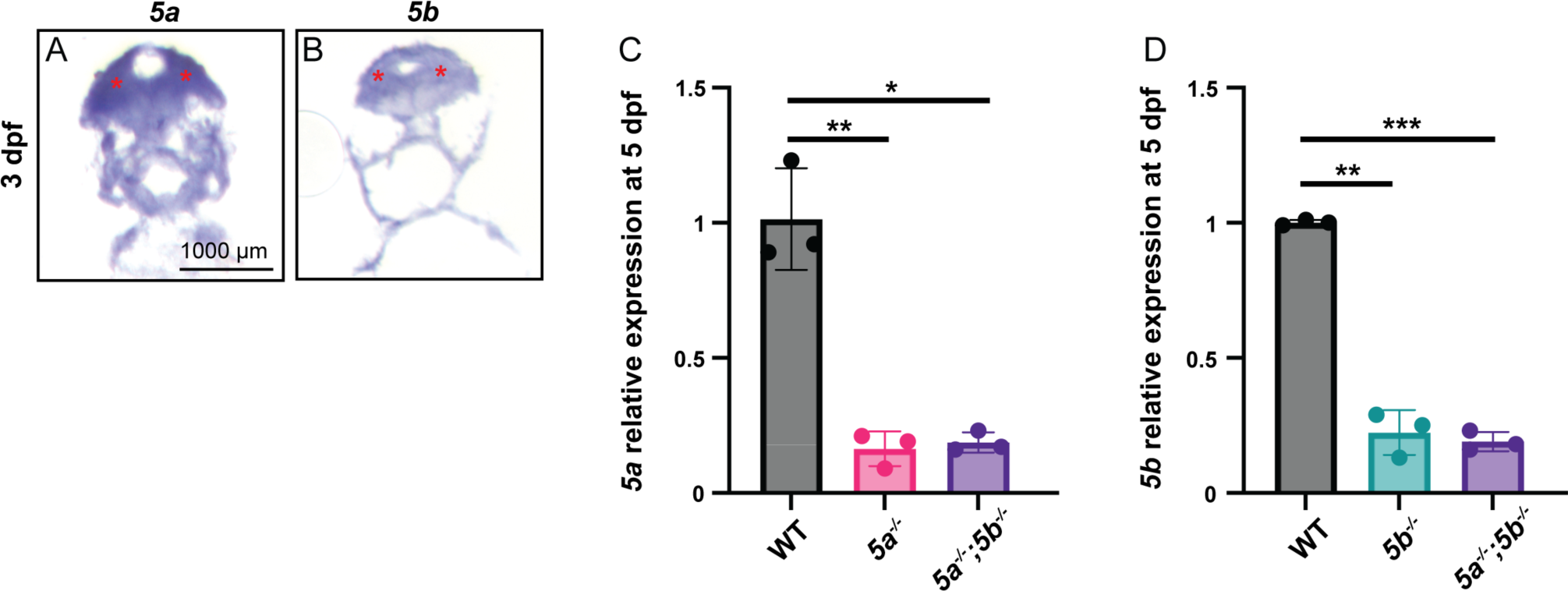
Expression analysis of *slc13a5* zebrafish paralogs (*5a* and *5b*) and transcript level detection in *slc13a5* mutants. (A, B) Sections of 3 dpf larvae post in situ hybridization for *5a* and *5b* expression. *5a* and *5b* expression is observed in several brain regions including the optic tectum region of MB, as indicated by red asterisks. (C) qPCR analysis for relative *5a* mRNA expression in 5 dpf *5a^−/−^* and *5a^−/−^;5b^−/−^ larvae* compared to WTs. WT, *5a^−/−^* and *5a^−/−^;5b^−/−^*, n = 3 × 10 larvae assessed as three biological and two technical replicates each. *5a* is downregulated in *5a^−/−^* and *5a^−/−^;5b^−/−^*larvae compared to WTs, indicating active mRNA degradation of *5a^−/−^* transcripts (D) qPCR analysis for relative *5b* mRNA expression in 5 dpf *5b^−/−^* and *5a^−/−^;5b^−/−^* compared to WTs. WT, *5b^−/−^ and 5a^−/−^;5b^−/−^*, n = 3 × 10 larvae assessed as three biological and two technical replicates each. *5b* is downregulated in *5b^−/−^ and 5a^−/−^;5b^−/−^*larvae compared to WTs, indicating active mRNA degradation of *5b^−/−^* transcripts. Data are Mean ± S.D., *P ≤ 0.05, **P ≤ 0.01, ***P ≤ 0.001-Unpaired t-test.

**Figure S2.**
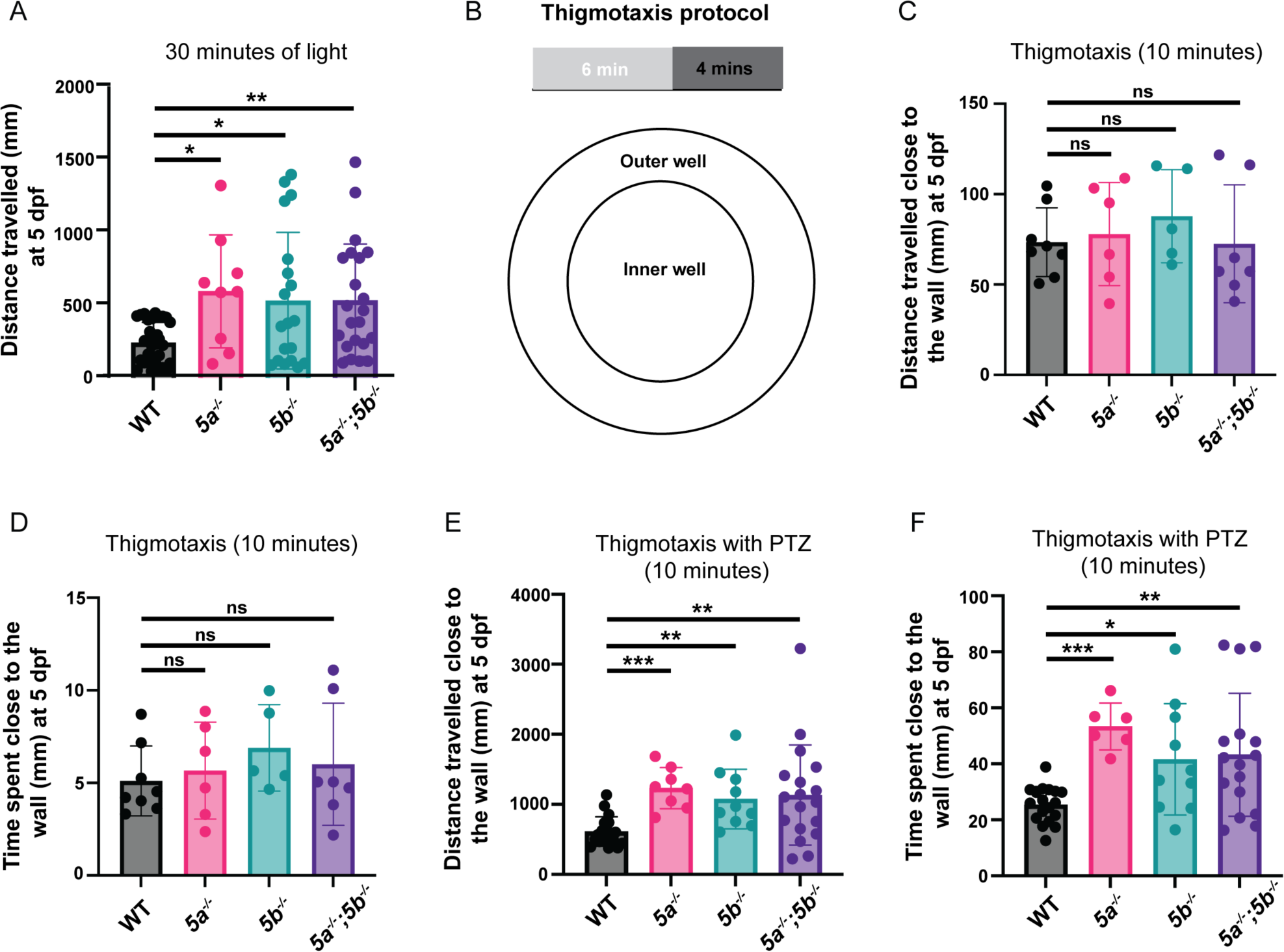
Analysis of spontaneous behavior and thigmotaxis in *slc13a5* mutants. (A) Quantification of distance travelled in 100% light for 30 minutes at 5 dpf. *slc13a5* mutants move higher distances compared to WTs during daytime. WT, n=33; *5a^−/−^,* n=9; *5b^−/−^*, n=19; *5a^−/−^;5b^−/−^*, n=22. (B) Schematic representation of thigmotaxis protocol. (C) Quantification of distance travelled close to the wall in 10 minutes. No significant changes were observed in the distance travelled between *slc13a5* mutants and WTs. WT, n=8; *5a^−/−^,* n=6; *5b^−/−^*, n=5; *5a^−/−^;5b^−/−^*, n=7. (D) Quantification of time spent close to the wall in 10 minutes. No significant changes were observed in the time spent between *slc13a5* mutants and WTs. WT, n=8; *5a^−/−^,* n=6; *5b^−/−^*, n=5; *5a^−/−^;5b^−/−^*, n=7. (E) Quantification of distance travelled close to the wall in 10 minutes post PTZ exposure. *slc13a5* mutants swam significantly higher distance hugging the wall compared to WT. WT, n=18; *5a^−/−^,* n=8; *5b^−/−^*, n=10; *5a^−/−^;5b^−/−^*, n=18. (F) Quantification of time spent close to the wall in 10 minutes post PTZ exposure. *slc13a5* mutants spent significantly more time hugging the wall compared to WT. WT, n=18; *5a^−/−^,* n=6; *5b^−/−^*, n=10; *5a^−/−^;5b^−/−^*, n=16. Data are Mean ± S.D., ns: no significant changes observed, *P ≤ 0.05, **P ≤ 0.01, ***P ≤ 0.001-Unpaired t-test. PTZ, pentylenetetrazol.

**Figure S3.**
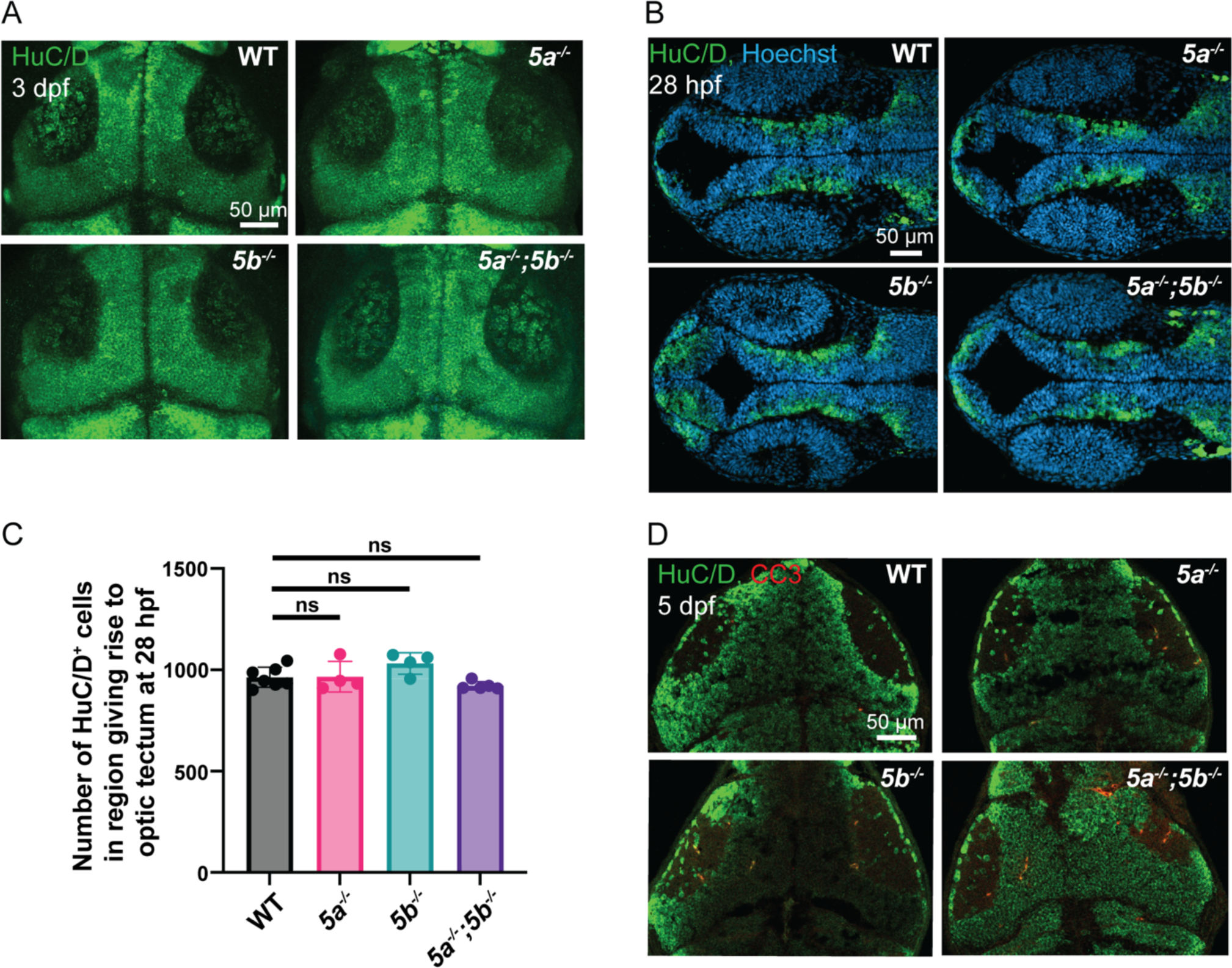
Neuron population and apoptosis analysis in *slc13a5* mutants. (A) 3 dpf WTs and *slc13a5* mutants; α-HuC/D (green). (B) 28 hpf WTs and *slc13a5* mutants; α-HuC/D (green), Hoechst (blue). (C) Quantification of HuC/D^+^ cells in the region that gives rise to optic tectum at 28 hpf. HuC/D^+^ cell numbers are unaffected in the *slc13a5* mutants compared to WTs. WT, n=7; *5a^−/−^,* n=4; *5b^−/−^*, n=4; *5a^−/−^;5b^−/−^*, n=5. (D) 5 dpf WTs and *slc13a5* mutants; α-HuC/D (green), CC3 (red). Data are Mean ± S.D., ns: no significant changes observed-Unpaired t-test.

**Figure S4.**
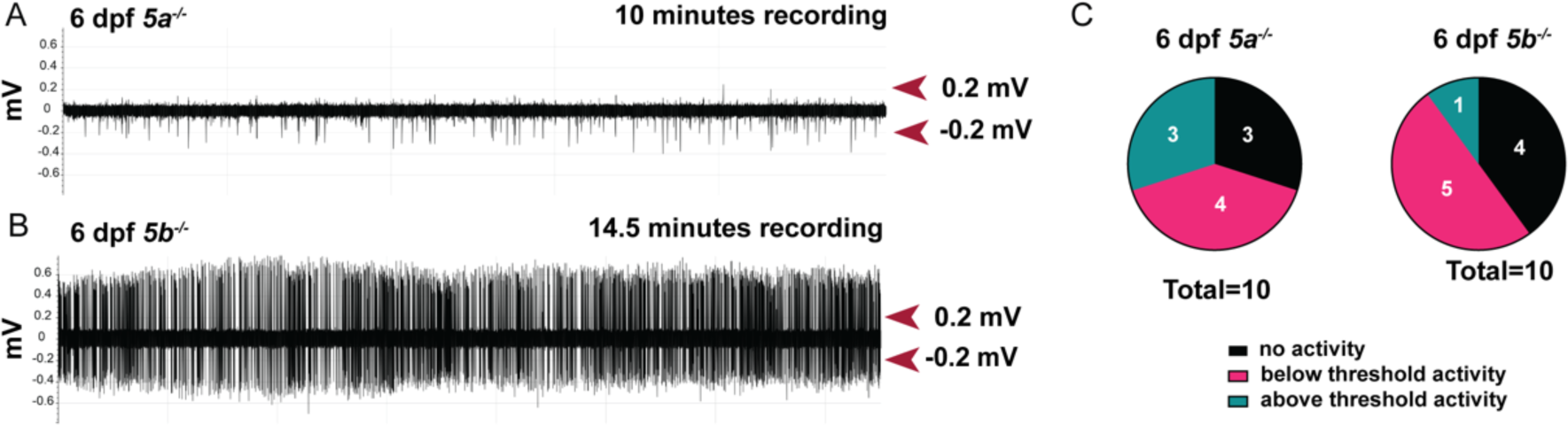
Electrophysiological analysis in *slc13a5* mutants. (A-C) Representative extracellular recordings obtained from optic tectum of 6 dpf *slc13a5* mutants, and a pie chart of number of *slc13a5* mutants showing different patterns of activity. The repetitive inter-ictal like discharges (<1s duration) with above threshold (>0.2mV), high-frequency, large-amplitude spikes seen in *slc13a5* mutants are indicative of increased network hyperexcitability. *slc13a5^−/−^* larvae also displayed below threshold (<0.2mV) brain activity.

**Figure S5.**
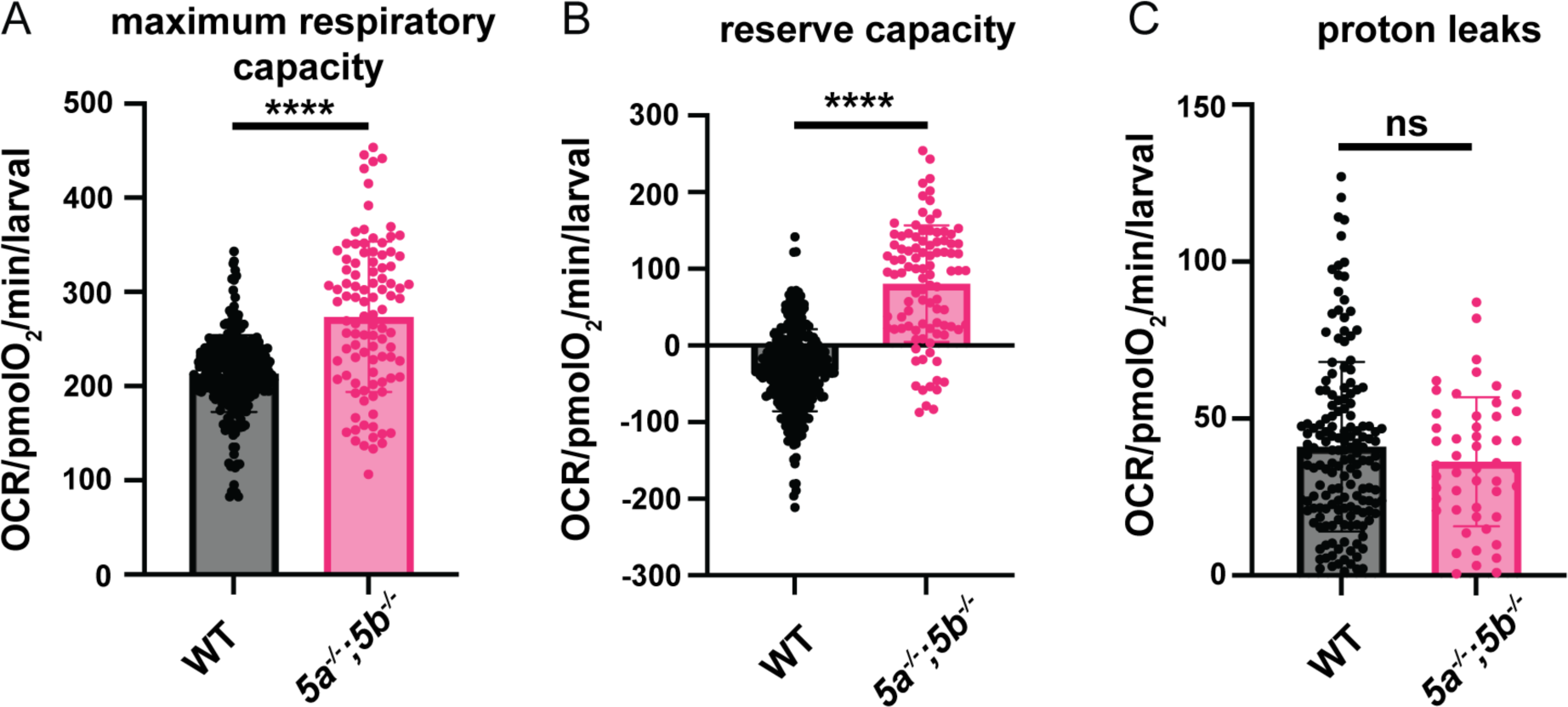
Bioenergetics profiling of *slc13a5* mutants. (A) Quantification of maximum respiratory capacity at 6 dpf. *slc13a5* mutants exhibit a significant increase in maximum respiratory capacity compared to WT. WT, n=58; *slc13a5* mutants, n=20 (individual values plotted from five cycles). (B) Quantification of reserve capacity at 6 dpf. *slc13a5* mutants exhibit a significant increase in reserve capacity compared to WT. WT, n=58; *slc13a5* mutants, n=20 (individual values plotted from five cycles). (C) Quantification of proton leaks at 6 dpf. This parameter was unchanged in *slc13a5* mutants compared to WT. WT, n=58; *slc13a5* mutants, n=20 (individual values plotted from five cycles). Data are Mean ± S.D., ns: no significant changes observed, ****P ≤ 0.0001-Unpaired t-test.

**Figure S6.**
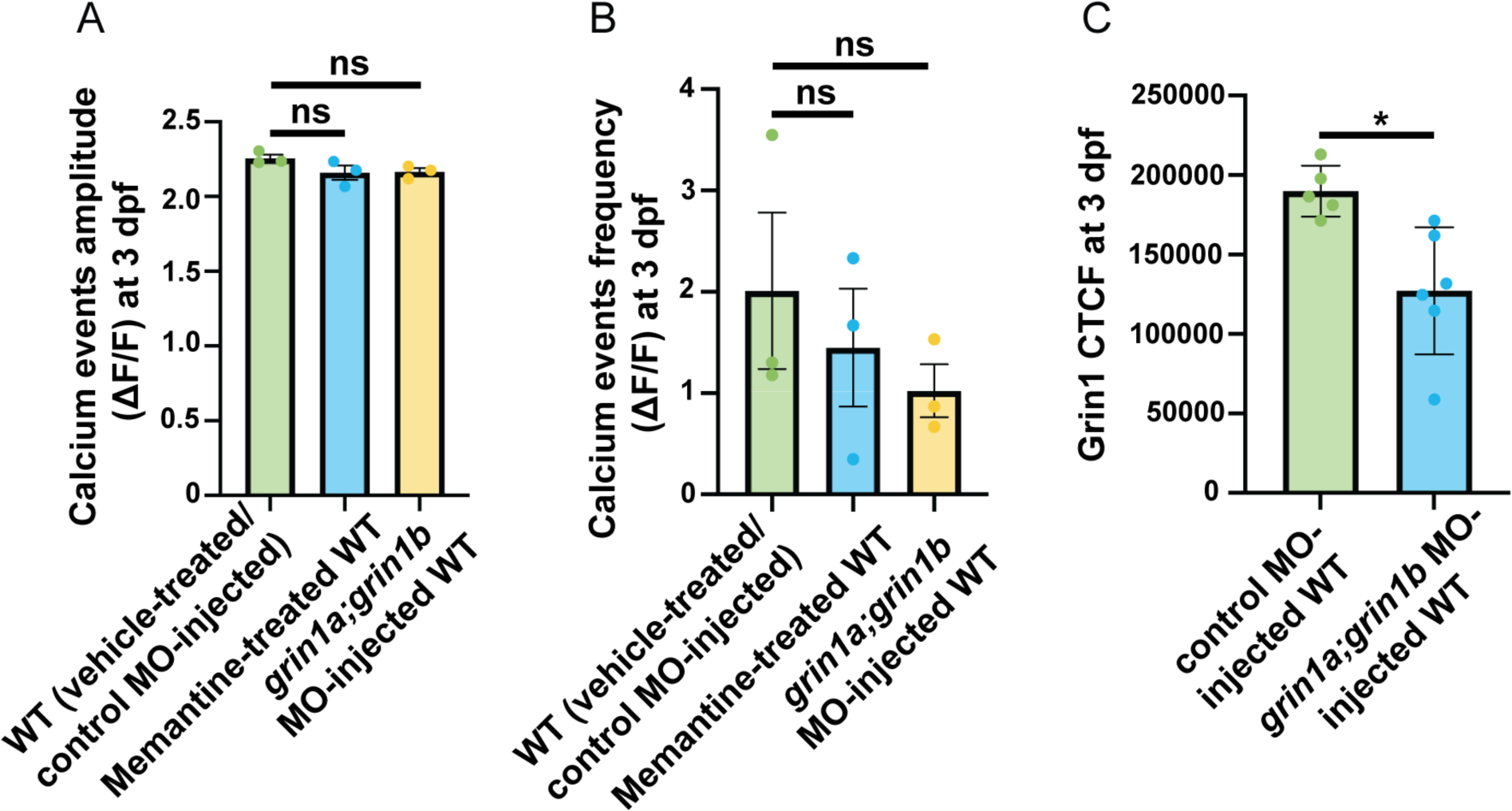
Assessment of the effect of suppressing NMDA receptor signaling on calcium influx in WTs and Grin1 expression after *grin1a;grin1b* MO injection. (A-B) Quantification of the amplitude and frequency of calcium events (ΔF/F>2) at 3 dpf. Memantine-treatment and *grin1a;grin1b* MO injections have no significant effect on the calcium events in WTs. Vehicle-treated/control MO-injected WT, n=5; memantine-treated WT, n=6; *grin1a;grin1b* MO injected WT, n=6. (C) Quantification of the fluorescence intensity (CTCF) of Grin1 (α-Grin1). *grin1a;grin1b* MO injections lead to significant reduction in the expression of Grin1 compared to control MO-injected WTs at 3 dpf. control MO-injected WT, n=5; *grin1a;grin1b* MO-injected WT, n=6. Data are Mean ± S.E.M. and Mean ± S.D., ns: no significant changes observed, *P ≤ 0.05-Unpaired t-test. CTCF, corrected total cell fluorescence.

**Figure S7.**
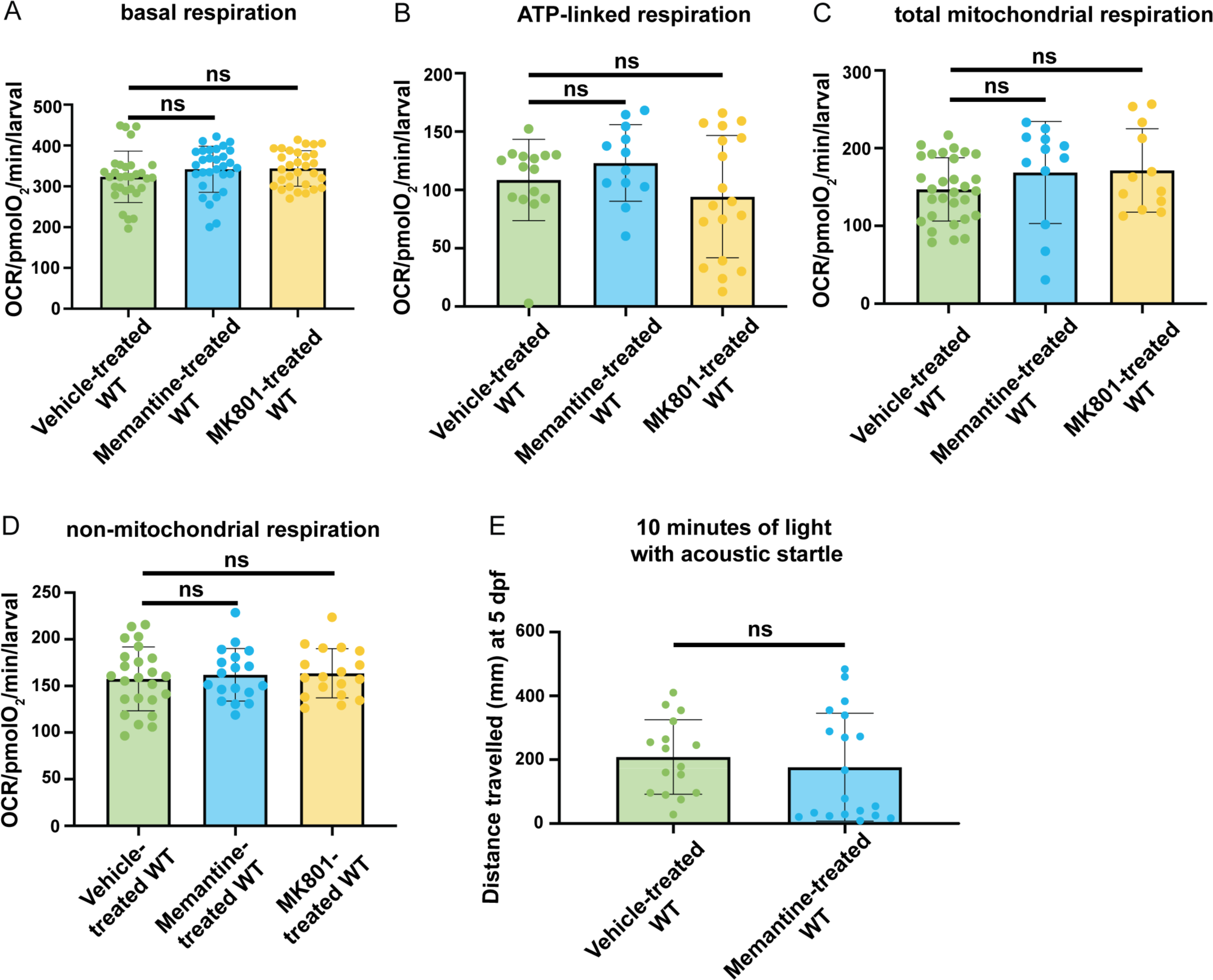
Effect of NMDA receptor antagonists on the bioenergetics parameters of WTs and on WT behavior under acoustic startle. (A) Quantification of basal respiration at 6 dpf. Memantine and MK801 treatments do not affect basal respiration in WTs. Vehicle-treated WT, n=6; memantine-treated WTs, n=6; MK801-treated WTs, n=6 (individual values plotted from five cycles). (B) Quantification of ATP-linked respiration at 6 dpf. Memantine and MK801 treatments do not affect ATP-linked respiration in WTs. Vehicle-treated WT, n=5; memantine-treated WTs, n=4; MK801-treated WTs, n=6 (individual values plotted from three cycles). (C) Quantification of total mitochondrial respiration at 6 dpf. Memantine and MK801 treatments do not affect total mitochondrial respiration in WTs. Vehicle-treated WT, n=10; memantine-treated WTs, n=4; MK801-treated WTs, n=4 (individual values plotted from three cycles). (D) Quantification of non-mitochondrial respiration at 6 dpf. Memantine and MK801 treatments do not affect non-mitochondrial respiration in WTs. Vehicle-treated WT, n=8; memantine-treated WTs, n=6; MK801-treated WTs, n=6 (individual values plotted from three cycles). (E) Quantification of total distance travelled in 10 minutes in acoustic startle before and after treatment with memantine at 5 dpf. Memantine treatment exhibited no effect on the behavior of WTs; Vehicle-treated WT, n=16; memantine-treated WT, n=19; MK801-treated WT, n=11. Data are Mean ± S.D., ns: no significant changes observed.

---

**Movie S1.** Movie (1000 fps) showing less calcium events in 3 dpf WT larvae in *Tg(elavl3:Hsa.H2B-GCaMP6s)* background.

**Movie S2.** Movie (1000 fps) showing more calcium events in 3 dpf *5a^−/−^;5b^−/−^* larvae in *Tg(elavl3:Hsa.H2B-GCaMP6s)* background.

**Movie S3.** Movie (1000 fps) showing more calcium events in 3 dpf vehicle-treated/control MO-injected *5a^−/−^;5b^−/−^*larvae in *Tg(elavl3:Hsa.H2B-GCaMP6s)* background.

**Movie S4.** Movie (1000 fps) showing less calcium events in 3 dpf memantine-treated *5a^−/−^ ;5b^−/−^* larvae in *Tg(elavl3:Hsa.H2B-GCaMP6s)* background.

**Movie S5.** Movie (1000 fps) showing less calcium events in 3 dpf *grin1a;grin1b* MO-injected *5a^−/−^;5b^−/−^* larvae in *Tg(elavl3:Hsa.H2B-GCaMP6s)* background.

**Table S1.**
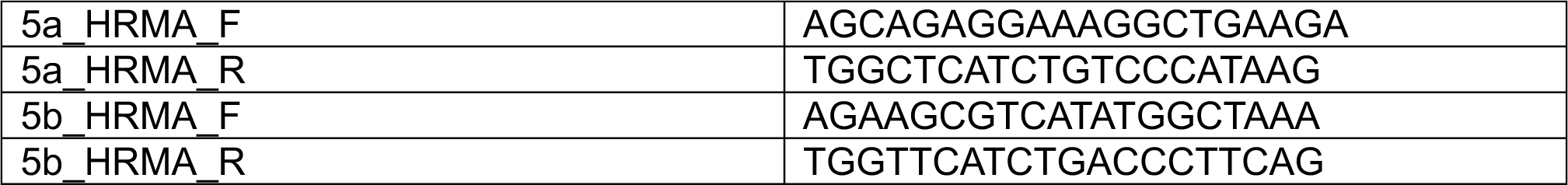
Primer list used for genotyping.

**Table S2.**
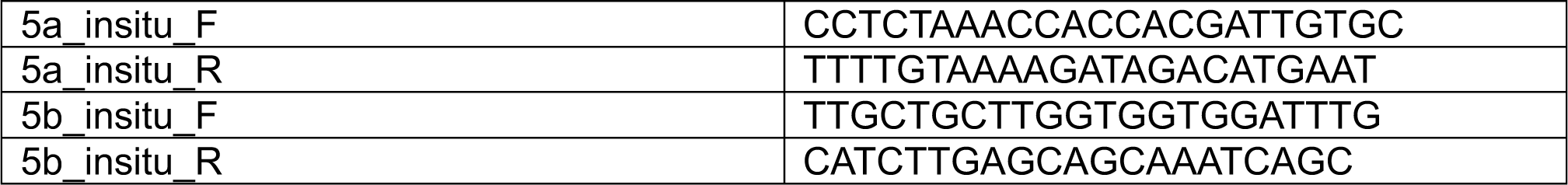
Primer list used to generate in situ hybridization probes.

**Table S3.**
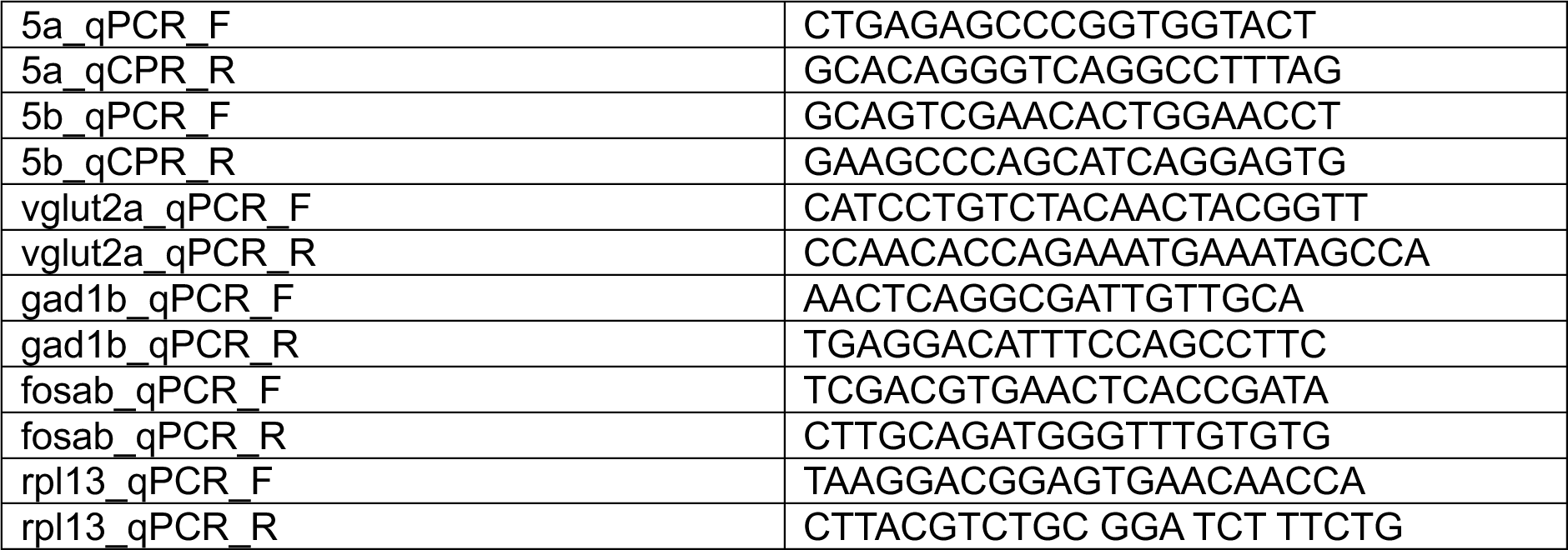
Primer list used in qPCR.

**Table S4.** Primary antibodies used in immunofluorescence.

| Primary antibody | Host species | Company | Catalog# | Concentration used |
| --- | --- | --- | --- | --- |
| Anti-HuC/D | Mouse | Invitrogen | A21271 | 1:200 |
| Anti-CC3 | Rabbit | BD Pharmingen | 559565 | 1:200 |
| Anti-Slc13a5 | Rabbit | Atlas Antibodies | HPA044343 | 1:100 |
| Anti-Gfap | Chicken | Abcam | ab4674 | 1:500 |
| Anti-pERK | Rabbit | CST | D13.14.4E | 1:200 |
| Anti-tERK | Mouse | CST | L34F12 | 1:200 |
| Anti-Grin1 | Mouse | Synaptic Systems | 114 011 | 1:500 |

## Notes

### Competing Interest Statement

The authors have declared no competing interest.

